# Widespread transcriptional memory shapes heritable states and functional heterogeneity in cancer and stem cells

**DOI:** 10.1101/2025.08.21.671653

**Authors:** Yongjie Lin, Xiangru Chen, Long Wu, Yutong Zhou, Yihan Lin

## Abstract

Recent studies show that non-genetic cellular heterogeneity, particularly through heritable cell states, fundamentally shapes cancer evolution and developmental trajectories. However, conventional single-cell transcriptomic snapshots lack temporal information needed to identify these heritable states. Here, we employ lineage-resolved single-cell transcriptomics to systematically map heritable cell states that persist across divisions, distinguishing them from transient fluctuations within a cell cycle. We uncover that heritable states are underpinned by widespread transcriptional memory, whereby gene expression is heritable, defining two classes of states in cancer and stem cells: clustered states, characterized by clustered gene expression, and latent states, marked by non-clustered gene expression. This memory shows unexpected conservation across cell types and conditions and is maintained by robust epigenetic mechanisms resistant to environmental perturbations. Functionally, memory genes predict critical behaviors including metastatic potential and lineage commitment, with latent-state genes often outperforming clustered-state genes. Our findings establish transcriptional memory as the molecular basis of heritable cellular heterogeneity, providing a framework for broadly understanding functional cellular variations across biological systems.

## Introduction

While it is well established that genetically homogeneous cell populations exhibit heterogeneous states^1–3^, the causes and functional roles of this heterogeneity remain largely unexplored^4–7^. Recent single-cell studies in cancer^8–11^ and developmental systems^12,13^ have demonstrated the biological importance of heritable cell states (**Figure 1A**). These states can be maintained across mitotic divisions and are thus clonally propagated, contrasting with transient states arising from cell cycle progression or stochastic gene expression fluctuations. For example, heritable states in melanoma cells characterized by high *NGFR* or *EGFR* expression can resist MEK inhibitor treatment *in vitro*^9^. In hematopoietic stem cells, heritable states associated with differential expression of transcription factors such as *Gata2* and *Gfi1* exhibit distinct lineage biases that are maintained across cell divisions, influencing long-term fate decisions during hematopoiesis^13^. Thus, it is becoming evident that decoding heritable cell states is necessary for understanding the significance of non-genetic heterogeneity. However, previous efforts have identified only a limited set of heritable cell states in a narrow range of systems, often regarding them as infrequent occurrences^8,10,11,13,14^. A comprehensive profiling is therefore essential to elucidate their ubiquity and roles. Existing lineage-tracing-based approaches to identify heritable cell states have constraints, limiting the depth and breadth of the investigation. Experimentally, to ensure the accurate and systematic capturing of heritable states, it would be necessary to densely sample a sufficient number of randomly seeded clones, but existing lineage-tracing approaches often optimize for multi-timepoint sampling^12^ and sparsely sample each clone^13^, or pre-sort cells to ensure defined initial states^11^ (Supplemental Note 1). Computationally, existing methods for processing lineage tracing data identify cell states as high-density cell clusters in low-dimensional embeddings of single-cell transcriptomic data^15–18^. However, such clustering approach cannot distinguish between transient gene expression dynamics (e.g., cell cycle-dependent dynamics) and stable, heritable gene expression differences, leading to an inaccurate picture of cellular heterogeneity^19^ (Supplemental Note 2). Consequently, for existing approaches, it is often necessary to first regress out the effect of known transient states (i.e., cell cycle states) before using clustering to identify cell states^20^, which could still be distorted by unknown transient states. Therefore, new approaches are needed to robustly resolve heritable cell states.

**Figure 1:**
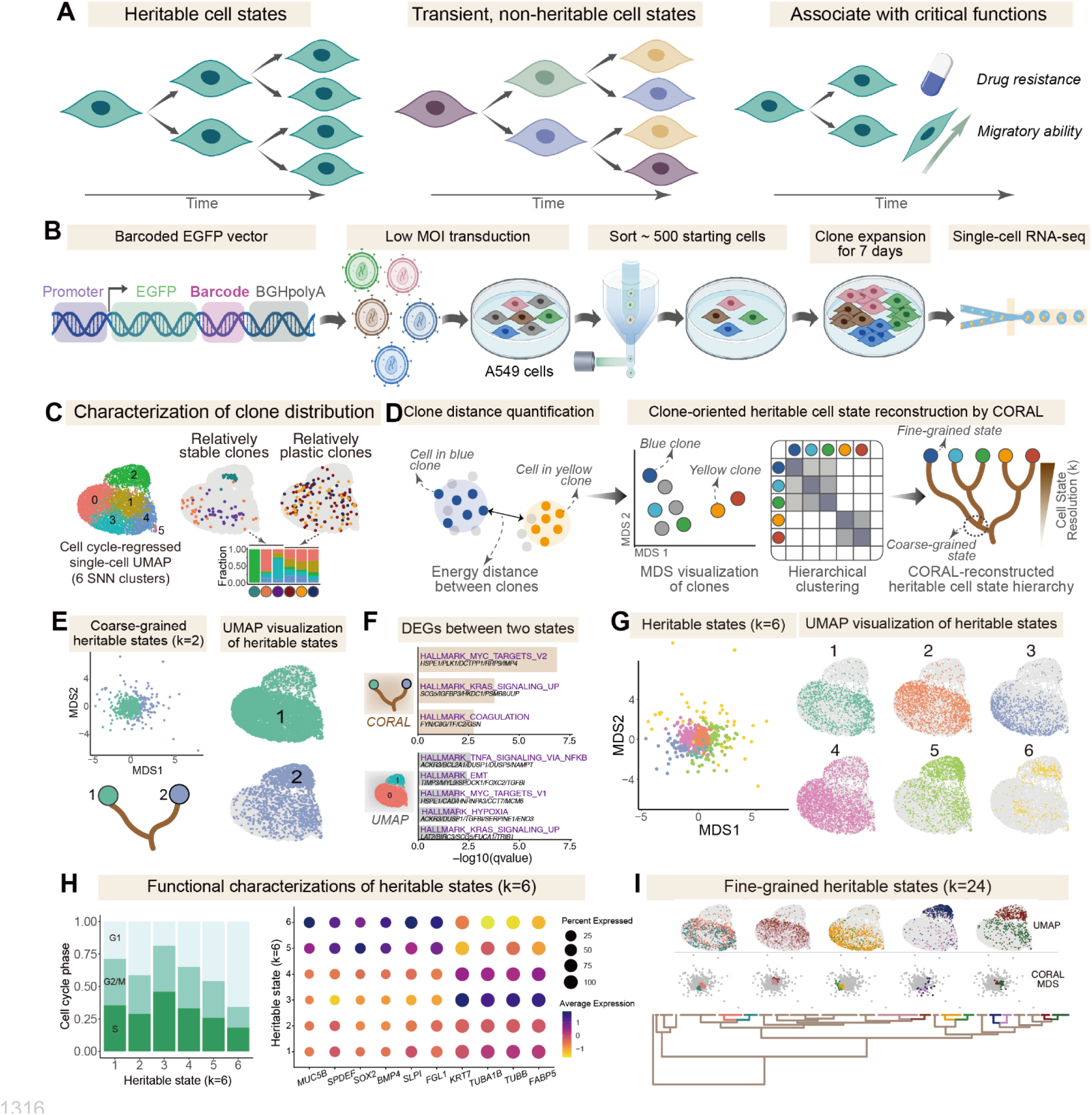
Lineage-resolved single-cell transcriptomics enables the reconstruction of a multi-scale heritable cell state hierarchy. **A**, Cartoons illustrating heritable states maintained across cell divisions (left), transient states (middle), and the associations of heritable states with critical functions (right). **B**, Design for lineage tracing experiment with dense clonal sampling using A549 lung cancer cells. Lentivirus library with EF1a promoter-*EGFP* and random barcodes in the 3’UTR was transfected into A549 cells. About 500 GFP+ transfected cells were sorted, plated, and cultured for 7 days. Single-cell RNA-seq was used to obtain transcriptomes of densely sampled clones. **C**, Clone-level behaviors of A549 cells. UMAP visualization of A549 cells (8972 cells) clustered into 6 SNN clusters (res=0.3) using cell-cycle regressed single-cell RNA-seq data (left). Certain clones were relatively localized to specific clusters, while some were relatively plastic, with cells spanning separate SNN clusters (right). **D**, Key steps in CORAL. Clone-clone similarities were quantified by energy distances, used for MDS visualization of clones. Hierarchical clustering defines heritable states at either coarse-grained or fine-grained resolutions, leading to a coral-like hierarchical cell state hierarchy. **E**, Two heritable states visualized on clonal MDS (left) and single-cell UMAP (right). Green state has 356 clones and 6558 cells, and blue state has 138 clones and 1908 cells. **F**, Gene Set Enrichment Analysis (GSEA) of differentially expressed genes (DEGs) between two CORAL-reconstructed heritable states (top) or between two states defined by conventional SNN clustering (bottom). **G**, Heritable states (k=6) visualized on clonal MDS (left) and single-cell UMAP (right). State 1: 50 clones, 1047 cells; state 2: 114 clone, 1686 cells; state 3: 50 clones, 1090 cells; state 4: 142 clones, 2735 cells; state 5: 95 clones, 1575 cells; state 6: 43 clones, 333 cells. **H**, Cell cycle phase proportion in each heritable state (left), and expression characteristics of representative state marker genes (k=6). Markers include goblet-like cell markers (*MUC5B*, *SPDEF*), stemness-related regulatory genes (*SOX2*, *BMP4*), senescence-associated genes (*SLPI*), cell cycle regulators (*TUBA1B*, *TUBB*), and potential cancer therapy targets (*FGL1*, *FABP5*). **I**, Fine-grained heritable states (k=24) visualized on a coral structure (bottom). Clones or cells from representative pairs of adjacent states were visualized clonal MDS (middle) or single-cell UMAP (top). Panels **a-b** were created with BioGDP.com^82^.

Here, we present an integrated experimental and computational workflow combining lineage-encoded single-cell transcriptomics with dense clonal sampling. We found that heritable states are prevalent in both cancer and developmental systems and across both *in vitro* and *in vivo* environments. One class of these states, clustered states, emerge as dominant clusters on single-cell UMAP, while the other class, latent states, appear generally diffusive on UMAP and are thus challenging to detect with conventional methods. We further uncovered pervasive transcriptional memory linked to these states, and revealed both clustered-state and latent-state memory genes. The memory property is unexpectedly conserved across cell types and environments, suggesting that it represents a gene-intrinsic property. Functionally, memory genes are enriched for core cellular functions and can predict critical cellular outcomes, including drug resistance, metastatic potential, and cell fate commitment. Intriguingly, latent-state genes appear to possess more predictive power than clustered-state genes, and the two memory gene classes are subjected to divergent epigenetic regulation. Collectively, this work establishes a framework for broadly and systematically decoding the mechanism and function of cellular heterogeneity in health and disease.

## Results

### An integrated experimental and computational workflow for profiling heritable cell states

Because heritable states are retained in cells of the same lineage, we hypothesized that single-cell clonal transcriptomics, where multiple single cells from a sufficient number of lineages (clones) are simultaneously measured by single-cell RNA-seq (**Figure S1A**), would be ideal for systematically profiling heritable cell states.

To test this, we used A549 lung cancer cell line as a model system, notable for its high degree of functional heterogeneity^21^ and quasi-mesenchymal characteristics^22^. By introducing lentiviral barcodes (expressing EGFP) into A549, we sorted ∼500 barcoded cells (i.e., progenitors for needed number of clones), cultured them for seven days, and then performed single-cell RNA-seq (**Figure 1B**, **Methods**). We recovered 8,538 barcoded cells spanning 549 clones, whereby ∼90% of clones (i.e., 494 clones) had at least 3 cells (**Figure S1B**).

Using this dataset, we investigated whether cell states are heritable during A549 clone formation, and developed a method to systematically identify these states while distinguishing them from non-heritable fluctuations, such as those tied to the cell cycle.

We first found that cells from the same lineage are more transcriptomically similar than randomly paired cells (**Figure S1C**) (consistent with previous reports in cancer^23^ and normal cells^24^), implicating the presence of heritable cell states for A549. At the individual clone level, we observed cells from certain clones are dominantly enriched in specific clusters on UMAP^25^ (defined by shared-nearest-neighbor (SNN) clustering of single-cell transcriptomes^26^) (**Figure 1C**), indicating cell state stability and consistent with the presence of heritability. Of note, not all cell states are heritable because some clones displayed relatively high plasticity, with cells scattering across multiple UMAP clusters (**Figure 1C**).

Thus, to assess the stability or heritability of cell states, it is necessary to examine multiple cells from the same lineage, and conventional clustering analysis at the single-cell level (without lineage information) cannot suffice. We therefore asked whether and how incorporating lineage information into single-cell transcriptomes would allow the systematic identification of heritable cell states. More generally, to reconstruct the cellular state landscape (i.e., Waddington’s landscape^27^), we developed CORAL (<u>c</u>lone-<u>o</u>riented reconstruction of cell state <u>a</u>ttractors in the landscape), a clone-based framework (**Figures 1D**, **S1D**, **Methods**). In this framework, “valleys” represent stable, heritable cell states (attractors) occupied by clonal populations, where cells maintain their states across divisions. “Hills” or “ridges” denote the topological barriers between these valleys, encoding transition difficulty between states (see Supplemental Note 3 for the theoretical basis of CORAL). Operationally, we calculated energy distances^28^ between clones, embedded clones in the classical multidimensional scaling (MDS) projection^29^ at the clone level (instead of the UMAP visualization at the single cell level), and used hierarchical clustering^30^ to define cell states as clusters of clones (**Figure 1D**). Of note, while conventional methods cluster individual cells based on transcriptomic proximity^20^, CORAL clusters individual clones. This fundamental shift in the unit of analysis (from a single cell to a clonal population) mitigates the formation of arbitrary clusters at high resolution, as states are defined by the collective, heritable properties of clonal populations, not the snapshot profiles of individual cells. And because heritable cell states were defined by applying hierarchical clustering to merge similar clones in the MDS embeddings, the ensemble of heritable cell states can be represented by a multi-scale “coral-like” structure (**Figure 1D**). In this structure, roots of larger branches at the bottom of the coral represent coarse-grained cell states (i.e., large clusters of clones), while the tips of smaller branches represent fine-grained cell states (i.e., small clusters occupied by one or few clones).

We first tested CORAL in silico, where it can successfully uncover heritable and hidden cell states (**Figures S1E-G**; **Table S1**; **Methods**). We next applied the approach to the A549 *in vitro* dataset. By visualizing cell states of individual clones on the MDS embedding (**Figure 1E** left), we found that clones form diffusive clusters, indicative of a high degree of state heterogeneity. We then clustered clones into two heritable states (**Figure 1E** left), and visualized individual cells from these two states on the single-cell UMAP (**Figure 1E** right), whereby cells from the two states recapitulate the dominant transcriptomic clusters (i.e., the two blobs on UMAP). Importantly, differentially expressed genes between the two states are more significantly enriched for biological functions than states defined by conventional clustering of single-cell transcriptomes (**Figure 1F**), underscoring the biological relevance of the identified heritable states. Of note, the previous UMAP (**Figure 1E** right) was constructed by using single-cell transcriptomes with cell cycle effects being regressed out (i.e., a standard pre-processing step in conventional single-cell analysis workflow^20^), while no pre-processing step was used for the CORAL workflow (**Figure 1E** left). And when cell cycle effects were not regressed away, the conventional single-cell analysis can no longer resolve the two cell states accurately (**Figure S2A**).

These results suggest that conventional single-cell workflow for identifying cell state is vulnerable to cell cycle artifacts (as previously reported^19^), and in contrast, CORAL robustly delineates heritable cell states and distinguishes them from transient states stemming from cell cycle effects, thereby underscoring the robustness of the clone-oriented approach for investigating heritable cell states with potential functional importance.

We next focused on the potential functional roles of heritable cell states. Given the diffusiveness of the MDS embedding, we chose to partition clones into six clusters (i.e., cell state resolution k=6) (**Figures 1G**, **S2B**; **Table S2**). We observed that these heritable cell states display distinct properties. For instance, the fraction of cells in G1 varied among states (**Figure 1H** left): state 3 contained relatively few G1-phase cells, whereas state 6 contained many, implicating differential proliferation capacities. Additionally, marker enrichment analyses (**Figure 1H** right) showed that state 5 and 6 expressed a suite of goblet-cell-associated genes (*MUC5B*, *CFH*, *SPDEF*) implicated in airway protection and lung regeneration^31^, consistent with the notion that cancer cell plasticity may give rise to states reminiscent of normal developmental lineages^32^. Notably, CORAL also pinpointed key regulators of these states, such as *SOX2* (known for promoting lung cancer metastasis) and *BMP4* (known for inducing G1 arrest in cancer cells), which were enriched in state 5, and *BMP4* alone in state 6 (**Figure 1H** right, **Figure S2C** left, **Figure S2D**). Both states also expressed secretory proteins tied to tumor prognosis (*SLPI5*^33^, *FGL1*^34^) (**Figure 1H** right), indicative of heightened tumorigenicity or immune evasion. Meanwhile, state 3, featuring the smallest fraction of G1 cells and marked by *KRT7* (known for promoting proliferation) (**Figure S2C** right), showed elevated levels of tubulin-related genes and metabolic regulators like *FABP5* (**Figure 1H** right), a potential cancer therapeutic target^35^.

At a higher cell state resolution (k=24), CORAL successfully uncovered relatively rare heritable states distinguished by subtle transcriptomic differences (**Figure 1I**), and depicted possible transitions between heritable states (**Figure S2E**).

Collectively, CORAL unveils the heterogeneous and often hidden heritable states within a lung cancer cell population, some of which may have implications in tumor progression and therapy adaptation.

### CORAL decodes functional heterogeneity in cancer and developmental systems

We next asked if clone-oriented cell state analysis approach would be broadly applicable. More specifically, we applied CORAL to another cancer model and a blood development model.

First, we tested CORAL on a lineage-tracing dataset of the WM989 melanoma cell line^11^, which captures transitions from drug-sensitive cells to “primed” drug-resistant states (**Figure 2A**; **Methods**). In this dataset, randomly selected clones and clones enriched for the primed *EGFR*+/*NT5E*+ phenotype were together profiled by single-cell RNA-seq (**Figures S3A-B**). When performing CORAL analysis, we first replicated the main findings of the analysis from the literature^11^, namely that the *SOX10*+/*EGFR*-(drug-sensitive) and *EGFR*+/*NT5E*+ (primed, drug-resistant) states segregate both in UMAP embedding (single-cell level) and MDS embedding (clone level), and that transitions can occur between these two states (**Figure 2B**). Critically, we further discovered an additional axis of heterogeneity—beyond the known *EGFR* axis— that divided the melanoma clones into four dominant heritable states (**Figure 2B**). Specifically, *ARF5* (ADP-ribosylation factor 5) was identified as another major determinant of clonal differences, while its expression pattern remained obscured in single-cell UMAP (which would not be possibly identified using conventional single-cell analysis) (**Figure S3B**). Intriguingly, ARF family proteins have been broadly implicated in tumorigenesis^36–39^, including tumor proliferation and invasiveness, and *ARF5* has been linked to melanoma progression^40^.

**Figure 2:**
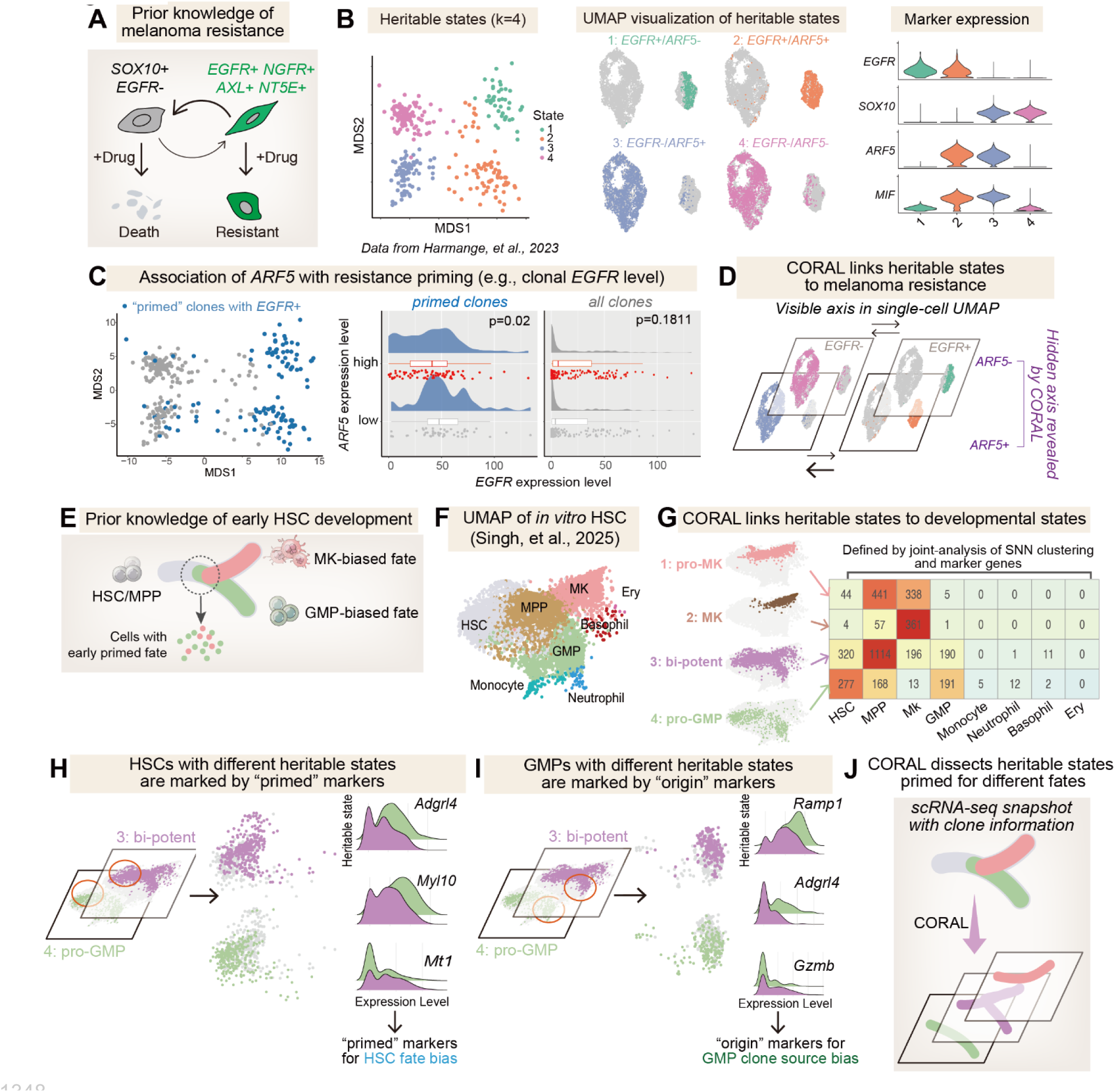
CORAL reveals heritable states linked to important functions in both cancer and developmental systems. **A-D**, CORAL revealed heritable states associated with melanoma drug resistance using public lineage tracing data of melanoma cell line WM989^11^. **A**, Known biology of the *EGFR*-relevant drug-resistance model in melanoma cells, where cells stochastically transition between a *SOX10*+/*EGFR*-sensitive state and a rare *EGFR*+/*NT5E*+ resistant state^11^. **B**, Heritable states (k=4) visualized on clonal MDS (left) and single-cell UMAP (middle), and expression patterns of key marker genes (right). State 1 (green, *EGFR*+/*ARF5*-): 45 clones, 563 cells; state 2 (orange, *EGFR*+/*ARF5*+): 75 clones, 1332 cells; state 3 (blue, *EGFR*-/*ARF5*+): 90 clones, 2926 cells; state 4 (magenta, *EGFR*-/*ARF5*-): 111 clones, 2760 cells. **C**, Association of *ARF5* level with resistance priming. Within the 122 “primed” clones with initial *EGFR*-high state (blue), *ARF5*-high clones tend to transition to *EGFR*-low state. **D**, CORAL elucidates heritable states underlying melanoma drug resistance, whereby clonal ARF5 levels associate with a faster transition from *EGFR*+ to *EGFR*-. **E-J**, CORAL deciphered heritable states primed for different fates during early development of hematopoietic stem cell (HSC) using public lineaging tracing data of mouse HSC^43^. **E**, Prior biological knowledge of the early development process of HSC, which may differentiate into granulocyte-monocyte progenitors (GMP fate) or megakaryocyte progenitors (MK fate). **F**, UMAP visualization of 12,672 cells after *in vitro* HSC expansion for 7 days^43^, illustrating the structure of the early HSC development process. **G**, Four heritable states associated with distinct cell fate preferences were identified: pro-MK state (167 clones, 828 cells), MK state (94 clones, 423 cells), bi-potent state (397 clones, 1832 cells), and pro-GMP state (183 clones, 668 cells). **H**, Feature plots and ridgeline plots of representative differentially expressed genes in HSC cells from heritable states primed for bi-potent state (state 3) and pro-GMP state (state 4). **I**, Feature plots and ridgeline plots of representative differentially expressed genes in GMP cells from heritable states primed for bi-potent state (state 3) and pro-GMP state (state 4). **J**, CORAL successfully partitioned HSC clones into distinct heritable states primed with different developmental potentials.

To probe the functional significance of the newly defined *ARF5* axis, we focused on clones enriched for the primed *EGFR*+/*NT5E*+ phenotype (**Figure 2C**). Although *EGFR* and *ARF5* expression levels showed a slight positive correlation (**Figure S3C**), *EGFR*+ clones with high *ARF5* levels tended to lose *EGFR* over time (**Figure 2C**), suggesting that *ARF5* may predict drug-resistance trajectories. Consistent with this observation, *ARF5* can also predict the trajectories of other known resistance markers, including *AXL* and *NT5E* (**Figure S3D**). Thus, the newly identified *ARF5* heterogeneity axis may modulate transition kinetics within the drug-resistance landscape in melanoma.

Another intriguing marker gene for heritable states 3 and 5 is *MIF* (macrophage migration inhibitory factor), known to be overexpressed in many tumor cells including melanoma^41,42^ and was recently shown to associate with aggressive melanoma^40^. *MIF* is highly correlated with *ARF5* (**Figure S3D**) and both may together regulate tumor progression through the maintenance of heritable melanoma cell states. These findings underscore the power of CORAL to uncover hidden heritable states that have critical functional implications (**Figure 2D**).

Because stem cells are also highly heterogeneous^4^, we hypothesized that CORAL may be valuable for understanding developmental heterogeneity. We thus applied CORAL to a mouse hematopoietic stem cell (HSC) differentiation dataset^43^. In *ex vivo* cultures^44^, most HSC-derived clones initially transition into multipotent progenitors (MPPs) before adopting one of two dominant developmental paths: granulocyte-monocyte progenitors (GMPs) or megakaryocyte progenitors (MKs)^43^ (**Figure 2E**). Notably, some clones commit almost exclusively to one fate, potentially reflecting early “priming events” that occur without substantial transcriptomic reprogramming^13^. Because CORAL excels at detecting subtle transcriptomic differences predictive of cellular outcomes, we applied it to (public) clonally resolved snapshot data of differentiating HSCs at day 7 (i.e., single-timepoint)^43^ to characterize the cell state landscape and uncover evidence for early priming (**Figure 2F**).

We successfully partitioned HSC clones into four well-documented developmental states (**Figures S3E, 2G**), including early MK fate (pro-MK) containing MPPs and MKs, late MK fate predominantly MKs, bi-potent fate encompassing MPPs, MKs, and GMPs, and pro-GMP fate containing MPPs and GMPs. Critically, the successful identification of these states did not depend on common single-cell RNA-seq preprocessing steps, such as feature selection or removal of cell-cycle and ribosome-mediated effects (which were implemented in the literature where the dataset was from^43^).

We next explored whether our clone-oriented approach could resolve finer structures within classically defined cell types, specifically focusing on HSC fate (**Figure 2F**). Despite their seemingly similar transcriptomes in standard analyses, we found that HSCs assigned to bi-potent and pro-GMP cell states expressed distinct gene sets, some of which have been reported as markers of “fate preference” ^13^ (**Figures 2H**, **S3F-G**; **Table S3**). These observations support the idea that priming occurs in a handful of genes before major transcriptomic shifts. We further investigated how a clone’s “origin” identity affects gene expression post-differentiation. For instance, GMPs originated from bi-potent clones highly expressed *Gzmb* (granzyme B, a secreted protease important for cell killing), whereas GMPs originated from pro-GMP clones showed higher levels of *Ramp1* (receptor activity-modifying protein 1) and *Adgrl4* (adhesion G protein-coupled receptor l4) (**Figures 2I**, **S3F-G**). Similar patterns were observed in MKs (**Figures S3H**, **S3F-G**), where early-MK, late-MK, and bi-potent clones each exhibited characteristic expression signatures.

Notably, the newly identified primed or origin markers only partially overlapped with classical cell-type markers defined by single-cell transcriptome data alone (**Figure S3I**). Nonetheless, these heritable state markers align with clone trajectories observed in multi-sampled HSC lineage-tracing studies^13,17,43^, underscoring the capacity of CORAL to detect functional critical heritable blood cell state markers by only using single-timepoint snapshot single-cell data (containing lineage information) (**Figure 2J**).

Together, these results demonstrate that clone-oriented analysis can successfully uncover hidden but functionally important heritable states as well as associated marker genes in diverse contexts, including cancer and development.

### Clone-oriented analysis uncovered widespread transcriptional memory

Because cell states are defined by the expression of cell state-related genes, it is natural to hypothesize that heritable cell states are linked to heritable gene expression. We thus need to decode gene expression heritability in order to understand cell state heritability. The ability of gene expression to be inherited across cell divisions—a phenomenon known as transcriptional memory—has been established in several systems^9,13,45–47^. However, the true pervasiveness of this memory remains unknown, and methods to systematically link it to heritable cell states from single-cell transcriptomes remain lacking.

To investigate transcriptional memory using single-cell transcriptomes, it is critical to discern the timescale of gene expression dynamics using clonal single-cell data. As each progenitor cell expands into a clonal population, certain genes may maintain their expression levels relatively steadily, exhibiting memory-like behavior heritable across cell divisions^9,24,48^ (**Figure 3A**). Other genes, such as those governing cell cycle progression, can change rapidly on timescales shorter than a single generation (**Figure 3A**). Distinguishing genes with slow dynamics from ones with fast dynamics is thus essential for identifying memory genes. To address this, we performed an ANOVA test for each gene, using omega squared^49^ as a measure of how much expression variance is explained by clone identity (**Figures 3B-C**; **Methods**). This statistic should effectively differentiate heritable, slow dynamic genes from quickly changing genes, and does not depend on the degree of freedom (compared with F-statistics in previous ANOVA-based method^24^).

**Figure 3:**
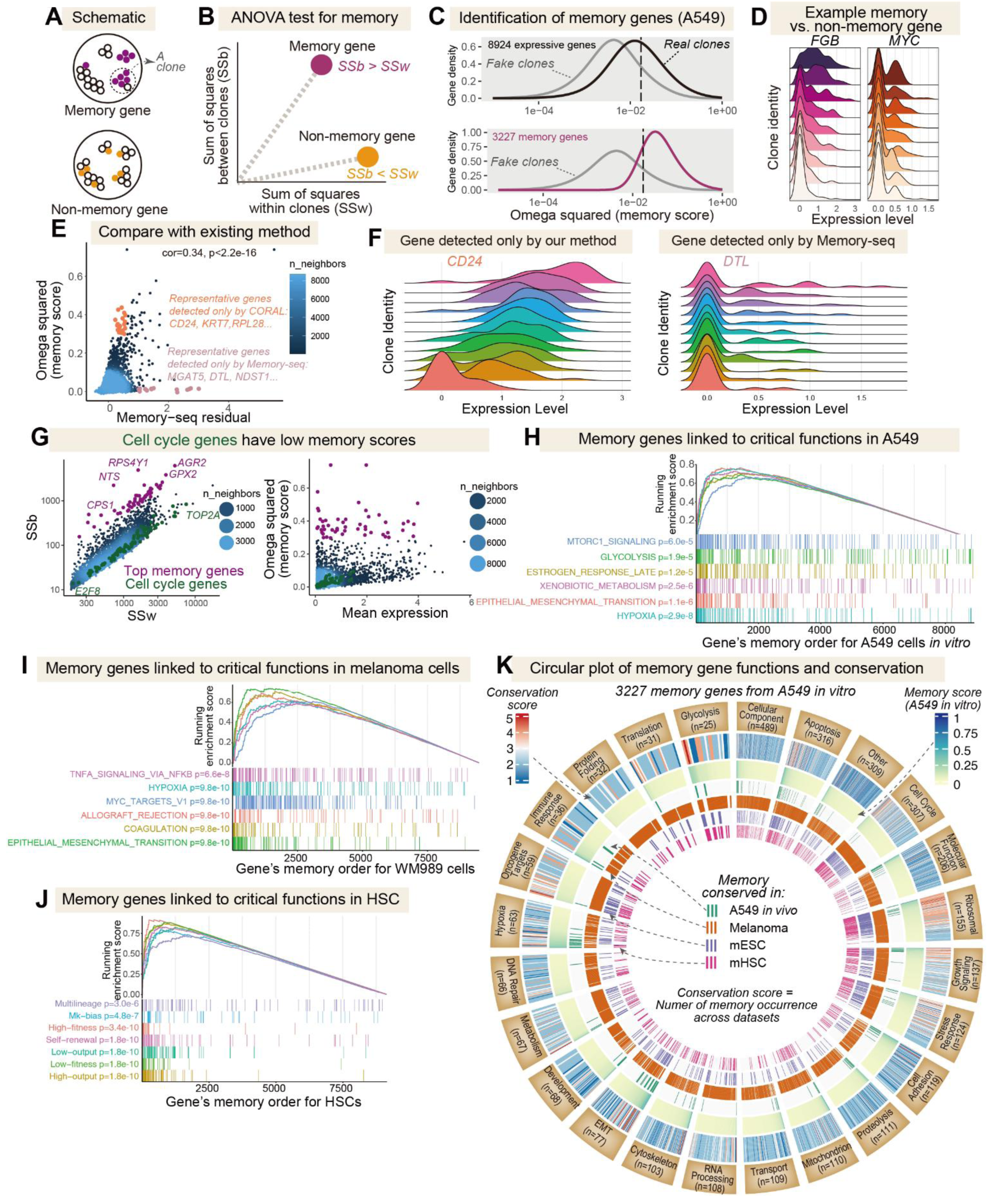
Clone-oriented analysis uncovers widespread and conserved transcriptional memory associated with key cancer and stem cell functions. **A**, Schematic of clonally heritable gene (top, i.e., memory gene) and non-heritable gene (bottom, i.e., non-memory gene). **B**, Memory genes exhibit higher inter-clone variance (quantified by SSb) than intra-clone variance (quantified by SSw), which can be detected by ANOVA test. **C**, Omega squared (an ANOVA test statistic, black) values of expressive genes (n=8924) for A549 cells *in vitro* compared with those of faked clones (gray) generated with randomly shuffled clone identity (top). 3227 genes (purple) were identified as memory genes (bottom). **D**, Clone-level expression distributions of example memory (*FGB*) or non-memory (*MYC*) gene. **E**, Comparison of our method and Memory-seq method for memory gene detection. While metrics of both methods for A549 dataset were correlated, significantly more memory genes can be detected by our method. **F**, Example false-negative (*CD24*) or false-positive (*DTL*) memory gene detected by Memory-seq method. **G**, Inter-clone (SSb) vs. intra-clone (SSw) variance (left), and omega squared statistic (right) of all expressed genes in A549 cells, with top 50 memory genes (purple) highlighted together with cell cycle genes (green). SSb: Sum of Squares between clones; SSw: Sum of Squares within clones. **H**, GSEA analysis using Hallmark gene sets for genes ranked by memory score for A549 *in vitro*. **I**, GSEA analysis using Hallmark gene sets for genes ranked by memory score for melanoma cells. **J**, GSEA analysis using HSC-critical gene sets for genes ranked by memory score for mouse hematopoietic stem cell (HSC). Gene sets were taken from the literature^43^. **K**, Functional landscape, and conservation of memory genes. Visualization of A549 *in vitro* memory genes (n=3227), their functional classifications, memory strength, and conservation across other four cellular contexts. Genes are grouped into sectors representing functional categories (including “Other” for unclassified genes, see **Methods**), with categories ordered by the number of genes they contain (n). Concentric tracks, from outermost to innermost, display: (1) functional category, (2) conservation score indicating the total number of datasets in which a gene’s memory status is conserved, (3) memory score (omega squared in **C**), and (4-7) four individual binary tracks denoting the conservation of memory.

Consistent with the presence of both slow and fast gene expression dynamics in A549 cells, we observed a broad distribution of omega-squared values (**Figure 3C**). To robustly separate these dynamics, we generated control (or fake) clones in which cell lineage information was randomly shuffled, thereby eliminating any genuine heritable patterns. As expected, the omega-squared values in these control clones were generally much lower than in real clones. By comparing these distributions, we identified more than 3,000 genes with significant clone-specific expression (i.e., memory genes) whose omega-squared values exceeded most of the omega-squared observed in control clones (**Figures 3C-D**; **Table S4**). Importantly, compared to Memory-seq^9^ (a reported method for identifying memory genes), our method more effectively identifies memory genes without an inherent bias for rare “ON” states (arising from the Memory-seq model assumption) (**Figures S4A**, **3E-F**), revealing that memory genes are far more pervasive than previously appreciated.

Specifically, we successfully segregated fast-changing cell-cycle genes from long-lasting memory genes, as reflected in their markedly different omega-squared values (**Figure 3G**). Among the memory genes, least-expected examples include ribosomal genes, traditionally viewed as housekeeping genes, emerged as highly heritable (**Figure S4B**). Consistently, recent studies suggest ribosomal gene expression can be clonally inherited in normal tissues^18,24^ and may decline with aging in single cells^50^, implicating epigenetic regulation over extended timescales. More importantly, we discovered that memory genes identified in A549 cells are broadly associated cancer-critical functions, including glycolysis, hypoxia, and epithelial-to-mesenchymal transition (EMT) (**Figure 3H**).

### Transcriptional memory associates with core cellular functions and is conserved across conditions and species

Given the surprising extent of transcriptional memory and its association with core functions, we sought to determine the robustness and pervasiveness of this phenomenon prior to exploring the linkage between memory genes and heritable cell states.

We first tested whether transcriptional memory and its functional association persist in A549 cells cultured *in vivo*. We thus performed clonal tracing experiment in an A549 cell-derived xenograft (CDX) model (**Figure S4C**), where barcoded cells were injected into mice and grown for 45 days (**Methods**). We applied CORAL to this *in vivo* dataset and partitioned clones into three heritable states (**Figure S4D**), whereby the cell state markers (**Figure S4E**) are distinct from those *in vitro* (**Figure 1H**), possibly reflecting different growth durations or microenvironmental cues. More importantly, for the 180 memory genes conserved in both *in vitro* (a total of 3227 memory genes, **Table S4**) and *in vivo* (a total of 252 memory genes, **Table S5**) datasets, their memory scores (quantified by omega squared statistic) were significantly correlated (**Figures S4F-G**). This observation is consistent with the picture that transcriptional memory is robust to culture conditions, and that memory genes in cancer cells are robustly associated with important functions (**Figures 3H, S4F**). Of note, the ability to detect memory genes using clone-oriented analysis can be limited by insufficient clonal sampling (i.e., not enough cells in a clone) or the extended clone formation time (leading to the loss of memory). And due to the very long period of *in vivo* tumor growth, a much lower number of genes was identified as memory genes compared to *in vitro* condition.

We next characterized transcriptional memory in the previous melanoma dataset^11^. Because melanoma cells were cultured for a duration comparable to cells in our A549 *in vitro* dataset, we would expect to identify pervasive memory genes if transcriptional memory is a gene-intrinsic feature and is thus conserved between cell types. Expectedly, we detected over 9000 memory genes in melanoma cells (**Table S6**), enriched for cancer-critical functions as in A549 (**Figure 3I**). Importantly, among these genes, 2593 of them overlapped with A549 cells and the memory scores were significantly correlated (**Figure S4H**). These results supported the hypothesis that transcriptional memory is a partially gene-intrinsic feature for human cancer cells and is highly associated with cancer-critical functions.

We next asked whether memory genes in mouse cells are also linked to key biological functions, and if the memory property is conserved across mouse cell types. For the mouse HSC dataset, we identified a total of 1512 memory genes (**Table S7**). Notably, 957 of them act as markers for different cell fates or different cell ancestries (**Table S3**), and these genes are enriched for critical functions in HSC (**Figure 3J**), including multi-lineage potential, high fitness, and output level control. To allow comparing with another mouse cell type, we applied CORAL to a public lineage-tracing murine embryonic stem cell (mESC) dataset^51^. After barcoding, mESC cells were expanded for 48 hours, providing only a short window for clone formation. Even with limited clonal sampling, we successfully identified heritable mESC cell states (**Figure S4I**). ANOVA analysis revealed 1044 memory genes (**Table S8**), enriched for critical functions relevant to self-renewal and differentiation (**Figure S4J**). By comparing memory genes between the two mouse cell types, we found 403 overlapping genes, whose memory scores were also correlated (**Figure S4K**) as observed for the human cell types.

To further explore the role and conservation of memory genes, we constructed a circular plot (**Figure 3K**; **Methods**) to illustrate the functional landscape and conservation of the 3227 memory genes identified from A549 *in vitro* dataset. These genes were broadly distributed across numerous functional categories, with notable groups including EMT, glycolysis, and stress response (**Figure 3K**, **Table S9**). A conservation score, ranging from 1 (A549 *in vitro* only) to 5 (conserved across all five analyzed datasets), showed diverse conservation patterns, with some genes broadly conserved while many others were context-specific. Notably, genes with high memory scores (positioned leftmost within each functional sector due to sorting) tend to exhibit correspondingly high conservation scores across diverse functional categories, a visual correlation observable by comparing the second and third tracks from the outermost edge (**Figure 3K**).

Collectively, these analyses revealed that transcriptional memory is an unexpectedly pervasive feature of gene expression. It is linked to core cellular functions, is robust to environmental changes, is conserved across human and mouse cells, and appears to be regulated in part by gene-intrinsic mechanisms.

### Memory genes are linked to heritable cell states in two distinct modes

Having analyzed memory genes and heritable cell states separately, we wondered how the heritability of memory genes is associated with heritable states, and what associative patterns exist. To address this, we first depicted how heritable states are organized, and then explored how such organization is underlined by memory genes.

Because the multi-scale coral-like structure of cell states (**Figure 1D**) is composed of coarse-grained and fine-grained heritable states, we wondered whether and how this hierarchy of heritable states can explain transcriptomic variance between clones. To do so, we calculated omega squared statistics from ANOVA tests across varying cell state resolutions using *in vitro* A549 dataset (**Figure 4A**). We found that while coarse-grained cell states only explained part of the inter-clone variance (**Figure 4A**), they captured the dominant heterogeneity visible in single-cell UMAP embeddings (**Figure 1E** right). Because these coarse-grained states appear as large clusters in single-cell UMAP (and can in principle be detected by conventional approach), we thus designated them as “clustered states”. As the resolution increases, increasingly more inter-clone variance was explained (**Figure 4A**), but the fine-grained states are challenging to identify with conventional approach because these states are not distinctively visible as large clusters on single-cell UMAP and are typically mixed up with other states (e.g., as in **Figure 1I**), which we thus designated as “latent states”. We therefore use clustered and latent states on a multi-scale coral-like structure to describe heritable cell state organization.

**Figure 4:**
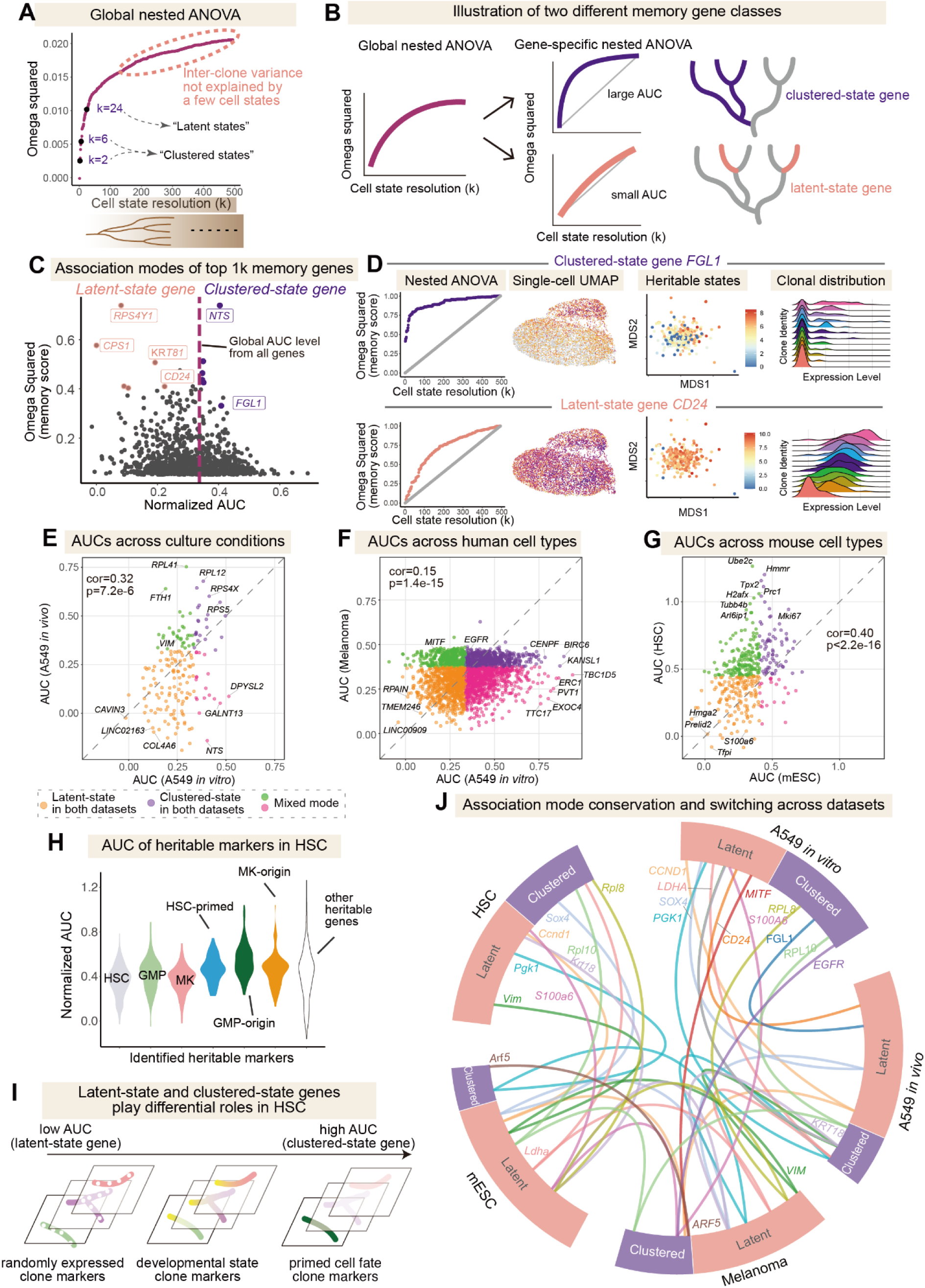
Memory genes define heritable states through two distinct and conserved modes of association. **A**, Variance explained (quantified by omega squared in global nested ANOVA test) by heritable states at different cell state resolutions. At low resolutions (e.g., k=2 or 6), coarse-grained states explained a large fraction of variance and captured dominant clusters on UMAP, designated as clustered states. At high resolution (e.g., k=24), fine-grained states cannot be distinctively visualized as clusters on UMAP and are thus designated as latent states. **B**, Gene-specific nested ANOVA analysis (left) could detect two modes of association between memory genes and heritable states (right). Clustered-state genes (purple) exhibit large area under the omega squared curve (AUC), while latent-state genes (orange) have small AUC. **C**, Quantification of association modes (i.e., AUC) for the top 1000 memory genes in A549 cells. Dashed line indicates the global area under the omega squared curve. **D**, Visualization of representative clustered-state (*FGL1*) and latent-state (*CD24*) genes, including the omega squared curve, single-cell UMAP, clonal MDS, and ridgeline plots of clonal gene expression distributions. **E**, Correlation analysis between A549 genes’ AUCs (i.e., association modes) *in vitro* and *in vivo*. n=180 shared memory genes. Color indicates whether and how the association mode is conserved between two datasets. **F**, Correlation between genes’ AUCs in A549 and melanoma cells. n=2593 shared memory genes. Same color scheme as **E**. **G**, Correlation between genes’ AUCs in mESC and mouse HSC cells. n=403 shared memory genes. Same color scheme as **E**. **H**, Distribution of AUCs for HSC memory genes identified as markers of cell type, primed events, original clone sources, or other (HSC: 346 genes; MK: 244 genes; GMP: 149 genes; HSC primed: 50 genes; GMP original: 73 genes; MK original: 95 genes; other memory genes: 555 genes). **I**, Latent-state and clustered-state genes play critical and differential roles in early HSC development. **J**, Conservation and switching of association modes of memory genes across five datasets, each represented by a sector of equal arc length. For each sector, genes are sorted by AUC values. Colored lines connect instances of the same gene across datasets. Genes described earlier as well as genes with highest conservation scores in Figure 3K are chosen for display.

We next focused on deciphering whether and how memory genes are associated with clustered or latent cell states. To do so, we needed to quantify how a memory gene’s variance between clones is explained by clustered or latent states. We proposed to compute each gene’s omega squared statistic across varying cell state resolutions (i.e., different numbers of cell states), producing a gene-specific curve (**Figure 4B**). The area under this curve (AUC) would thus capture how well a gene’s clonal variance is explained as cell state resolution increases. In theory, for genes with high AUC values, their clonal variances can be largely explained by coarse-grained or clustered cell states (**Figure 4B** top), because the omega-squared curve increased much faster at low resolution compared to at high resolution. In contrast, for genes with low AUC values, their clonal variances are better explained by fine-grained, latent cell states (**Figure 4B** bottom). Therefore, AUC metric would allow classifying the association pattern between memory genes and heritable cell states.

To test this, we calculated the AUC of the top 1,000 memory genes in the A549 *in vitro* dataset, revealing substantial variations in AUC values and thus their mode of association with heritable cell states (**Figure 4C**). High AUC genes such as *FGL1*^34^ (fibrinogen-like 1, a ligand of LAG-3 and a prognostic cancer marker) are linked to clustered cell states and would be effectively captured by dominant single-cell UMAP clusters. As expected, *FGL1* expression pattern forms a tight cluster on single-cell UMAP (**Figure 4D** top) and can in theory be detected by conventional single-cell analysis (i.e., analogous to conventional cell state markers). We therefore designated high AUC genes as “clustered-state genes”. In contrast, low AUC genes such as *CD24* (a well-known EMT and cancer stem cell marker) are linked to latent cell states, and formed a diffusive pattern on single-cell UMAP (**Figure 4D** bottom), which cannot be easily detected by conventional single-cell analysis (i.e., non-conventional, latent cell state markers) and are thus designated as “latent-state genes”. Of note, despite the distinct cell state associations, the two types of genes exhibited high expression correlation (**Figure S5A**). As a further note, the distinction between clustered-state and latent-state genes represents a continuum rather than a strict dichotomy, and intermediate genes may display characteristics of both modes.

We next compared AUC of memory genes between A549 cells *in vitro* and *in vivo*. We found that AUC’s are partially conserved (**Figures S5B, 4E**) and that genes can switch between high and low AUC states (**Figure 4E**). For example, *VIM*, a classical epithelial-mesenchymal transition (EMT) marker, maintained its memory property in both *in vitro* and *in vitro* A549 cells, but changed from a latent-state gene *in vitro* to a clustered-state gene *in vivo* (**Figure 4E**).

Across cancer cell types, we found that in melanoma cells, many memory genes displayed high AUC values and are thus more clustered-like (**Figures S5C**). As expected, clustered-state genes aligned primarily along the *EGFR* or *ARF5* axes (**Figure S5D**). While modes of association with heritable states (i.e., AUC) were partially conserved between A549 and melanoma cells (**Figure 4F**), it is less conserved than memory itself (**Figure S4H**). As a notable example, *MITF*, a prognostic marker in melanoma^52^, maintained its memory property in both lung cancer cells and melanoma cells, but changed from a latent-state gene in lung cancer cells to a clustered-state gene in melanoma cells (**Figure 4F**).

As for mouse cell types (mESC and HSC), we intriguingly found that more memory genes are latent-state genes rather than clustered-state genes (**Figure S5E-H**, **Tables S7-8**), and that the association mode is well-conserved for shared memory genes between mESC and HSC (**Figure 4G**). This suggests that a large fraction of memory genes that play critical roles in latent mouse cell states would not be easily identified with conventional approaches. Functionally notable latent-state genes in mESC (**Figure S5F**) include *S100a6* (a calcium binding protein), a known marker for differentiation trajectories^53^, and *Rian*, previously implicated in regulating the developmental potential of ESCs^54^ and iPSCs^55^. As such, while stem cells with distinct *Rian* levels would share similar transcriptome states (which cannot be easily distinguished by conventional analysis), ones with high *Rian* expression would mark a latent heritable state capable of full potency in embryo complementation assays^54^.

For HSC, newly identified primed and origin markers (a set of ∼200 genes) skewed toward higher AUC (**Figure 4H**), consistent with their robust association with distinct heritable cell states (i.e., clustered states). Conversely, traditional cell-type markers showed generally low to medium AUC (**Figure 4H**). These together painted a picture that latent-state and clustered-state genes play critical and differential roles in HSC development (**Figure 4I**).

Globally across all five datasets, we found that association mode of some orthologous genes can be conserved across species (**Figure 4J**). For example, *ARF5* and its ortholog *Arf5* are a clustered-state gene in human melanoma cells and mouse ESC respectively, while *VIM* (and its ortholog *Vim*) and *LDHA* (and its ortholog *Ldha*) are latent-state genes for both cell types (**Figure 4J**).

Together, using AUC metric we systematically associated memory genes to heritable cell states, and revealed two types of memory genes based on the association mode, whereby the association mode can be partially conserved between culture conditions and across cell types. High AUC, clustered-state memory genes act like conventional cell state markers, define major cell states and dominant UMAP clusters, and thus can be identified by conventional single-cell analysis. By contrast, low AUC, latent-state memory genes act as non-conventional and latent cell state markers, define latent cell states that are diffusive on UMAP, and thus cannot be easily identified by conventional single-cell analysis. More importantly, the discovery of latent-state genes as markers of previously hidden cell sates challenges the prevailing view that cell state markers should appear as clusters in single-cell embeddings^20,26,56^, which calls for further investigations of their functional implications and underlying mechanisms.

### Memory genes defining latent cell states predict EMT plasticity and drug resistance

In addition to earlier functional enrichment analyses (**Figures 3HJ, S4F, S4J**), we next sought to experimentally examine the predictive roles of transcriptional memory for core cellular functions, particularly genes linked to latent cell states. While the predictive roles of conventional cell state markers (e.g., clustered-state genes) have been widely recognized, the roles of non-conventional, latent cell state markers (e.g., latent-state genes) remained to be determined. Given the implication from earlier analysis (**Figure 3H**), we sought to experimentally test the hypothesis that latent-state genes might predict cancer-critical behaviors such as EMT phenotypes in A549 cells.

To this end, we designed phenotype-mapping assays to investigate whether and how pre-existing memory gene variation might predict cancer-critical behaviors (**Figure 5A**). Specifically, we used the parental population of the previous *in vitro* A549 experiment, comprising approximately 10,000 distinct clones (**Methods**). We subjected these cells to three enrichment protocols—gemcitabine treatment, Transwell migration, and 3D culture—then measured barcode representation to determine which clones were enriched in each condition (**Figure 5A**). Notably, clone enrichments across all three environments were positively correlated (**Figure 5B**), consistent with prior findings that cells exhibiting high EMT potential perform better under multiple stresses^57^.

**Figure 5:**
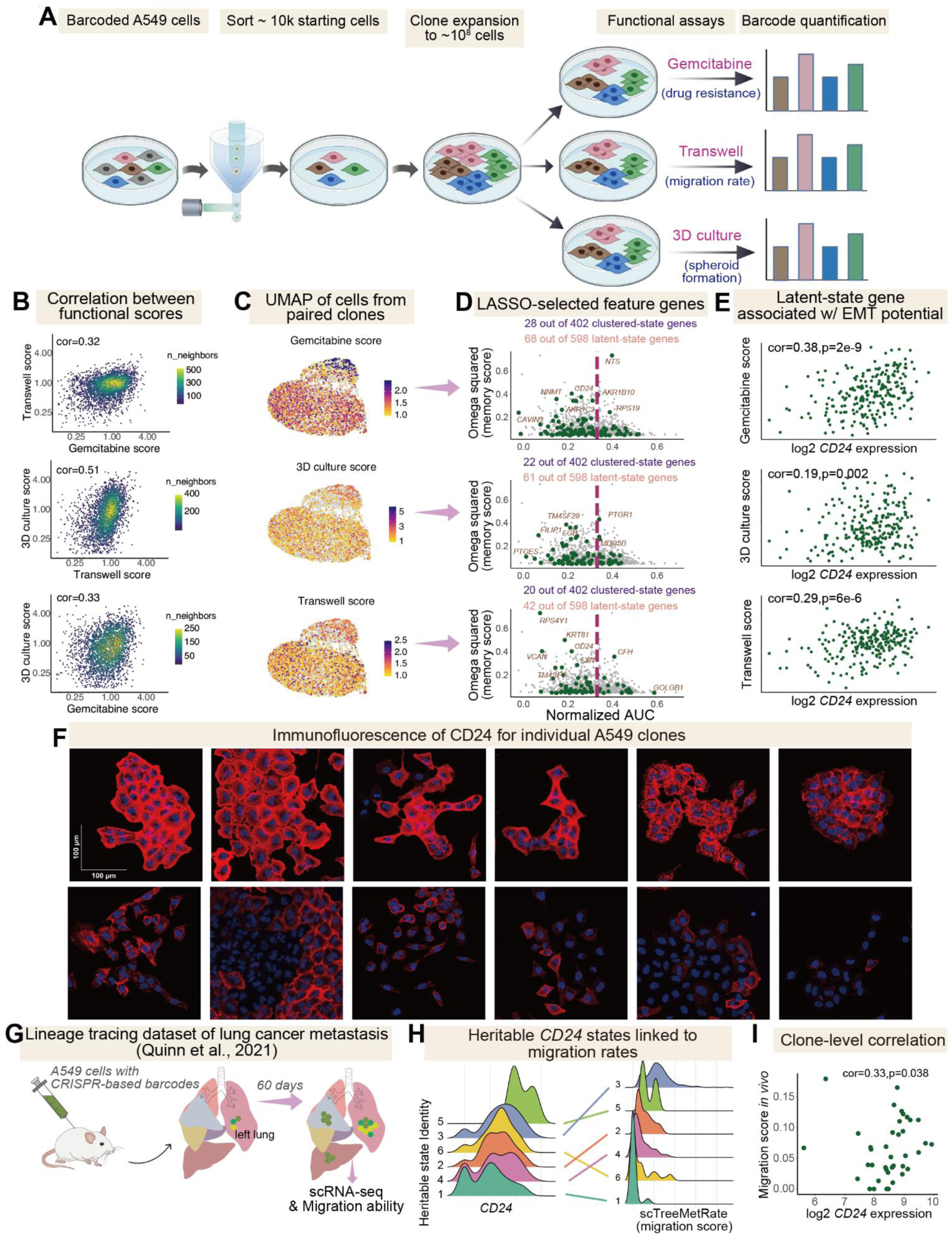
Memory genes associated with latent states possess higher predictive power for cancer-critical behaviors. **A**, Design of phenotype mapping experiment. The parental barcoded A549 population (with ∼10,000 uniquely labeled clones) was expanded to ∼10^8^ cells. Cells were then separately enriched through functional assays including gemcitabine treatment, Transwell migration, and 3D culture. Barcode abundance in each enriched population was quantified by NGS for further analysis. Created with BioGDP.com^82^. **B**, Correlation of functional enrichment scores. Density plot and Spearman correlation between functional enrichment scores (relative fold change compared to the control group) of different functional assays from the phenotype mapping experiment. n=3147 shared clones. **C**, Clones’ functional enrichment scores mapped to the single-cell UMAP, colored by functional enrichment scores at the clone level. Single-cell transcriptomes were measured in **Figure 1B**. **D**, Key feature genes (dark green) associated with clones’ functional scores were selected from top 1000 memory genes (gray) using LASSO. Note that more latent-state genes than clustered-state genes were selected by LASSO. **E**, Scatter plots showing the relationship between clone-level expression of LASSO-selected memory gene, *CD24*, and clonal functional scores for the largest 250 clones. Spearman correlation was indicated. **F**, Immunofluorescence images showing CD24 expression (red) for *in vitro* cultured A549 clones, sorted by the average expression level (from left to right, top to bottom). Cells were plated sparsely on culture plate and grown for 4 days. Blue indicates cell nuclei stained by DAPI. **G**, Overview of public lineage tracing dataset of lung cancer metastasis^73^. A549 cells labeled with CRISPR-based evolving barcodes were injected into the left lung of a mouse. After 60 days, samples from different tissues were collected for single-cell RNA-seq to measure both the transcriptome of cancer cells and their migration ability. **H**, Ridgeline plot of *CD24* expression and migration ability for different heritable states, showing that heritable *CD24* expression is linked to metastasis. **I**, Spearman correlation between *CD24* expression and migration ability at the clone level, calculated for the largest 50 clones.

To study the predictive roles of memory genes for these phenotypes, we mapped the functional enrichment scores to the clones whose single-cell transcriptomes were measured in our earlier experiment. We found that clone-level functional heterogeneity could not be explained by major cell clusters on UMAP (**Figure 5C, S6A**), implicating that the observed cancer-critical phenotypes do not arise from specific conventional cell states. This finding suggested that the observed phenotypes are not predicted by conventional cell states, but rather by the subtle variations defined by latent-state genes.

To test this, we employed LASSO regression^58^ to determine which memory genes best predict clonal functional phenotypes. Intriguingly, the top predictors predominantly consisted of latent-state genes rather than clustered-state genes (**Figures 5D, S6B**). Particularly, *CD24*, a latent-state gene, emerged as a strong explanatory variable and was positively correlated with the EMT-relevant phenotypes (**Figure 5E**). This aligns with the literature findings that certain epithelial markers, including *CD24*, can be linked to hybrid E/M phenotypes with elevated migratory and tumorigenic potential^59–61^. Our data suggest that in A549 cells, *CD24* may be associated with a continuum of EMT-relevant phenotypes at clone level across different clones (**Figure 4D** bottom), highlighting the critical role of non-conventional, latent cell state markers (which would be otherwise challenging to decipher with conventional methods).

Give the notable functional implications of latent-state genes, particularly *CD24*, we first validated that CD24 protein indeed exhibited clonal expression patterns in A549 similar to *CD24* mRNA (**Figure 5F** vs. **Figure 4D** bottom). We then asked whether it would also delineate important phenotypes *in vivo*. To investigate this, we examined a public lineage-tracing dataset that tracked metastatic progression of barcoded lung cancer cells in a xenograft model^62^ (**Figures 5G, S6C-D**). Intriguingly, heritable *CD24* states are associated with metastatic phenotype and clonal *CD24* expression level exhibited a significant correlation with migratory phenotype (**Figures 5H-I, S6E-F**), underscoring the capability of latent-state genes in predicting *in vivo* metastatic potential.

Together, these results provide support for the picture that latent-state genes can better predict cancer-critical functions than clustered-state genes, further underscoring the effectiveness of CORAL in decoding functional latent states.

### Epigenetic maintenance of memory under both normal and stress conditions

Thus far, we have focused on deciphering the heritability of gene memory during clone formation, which could persist over tens of cell divisions under normal conditions and predict critical functions at the clone level. However, it remains to be determined whether such memory, particularly the memory of latent-state genes, would also persist when cells are subjected to perturbations, which could have important implications for broadly understanding the functional roles of transcriptional memory.

To test this, we investigated whether cells with different baseline levels of the representative latent-state gene, *CD24*, might respond differently to external perturbations. We focused on TGF-β stimuli, a well-known inducer of EMT. While TGF-β is known to trigger substantial gene expression state transitions, including altered *CD24* expression^63^, less is known about whether heritable gene memory can be completely disrupted, or if the pre-stress gene expression state is maintained even after the withdrawal of the external cue (**Figure 6A**).

**Figure 6:**
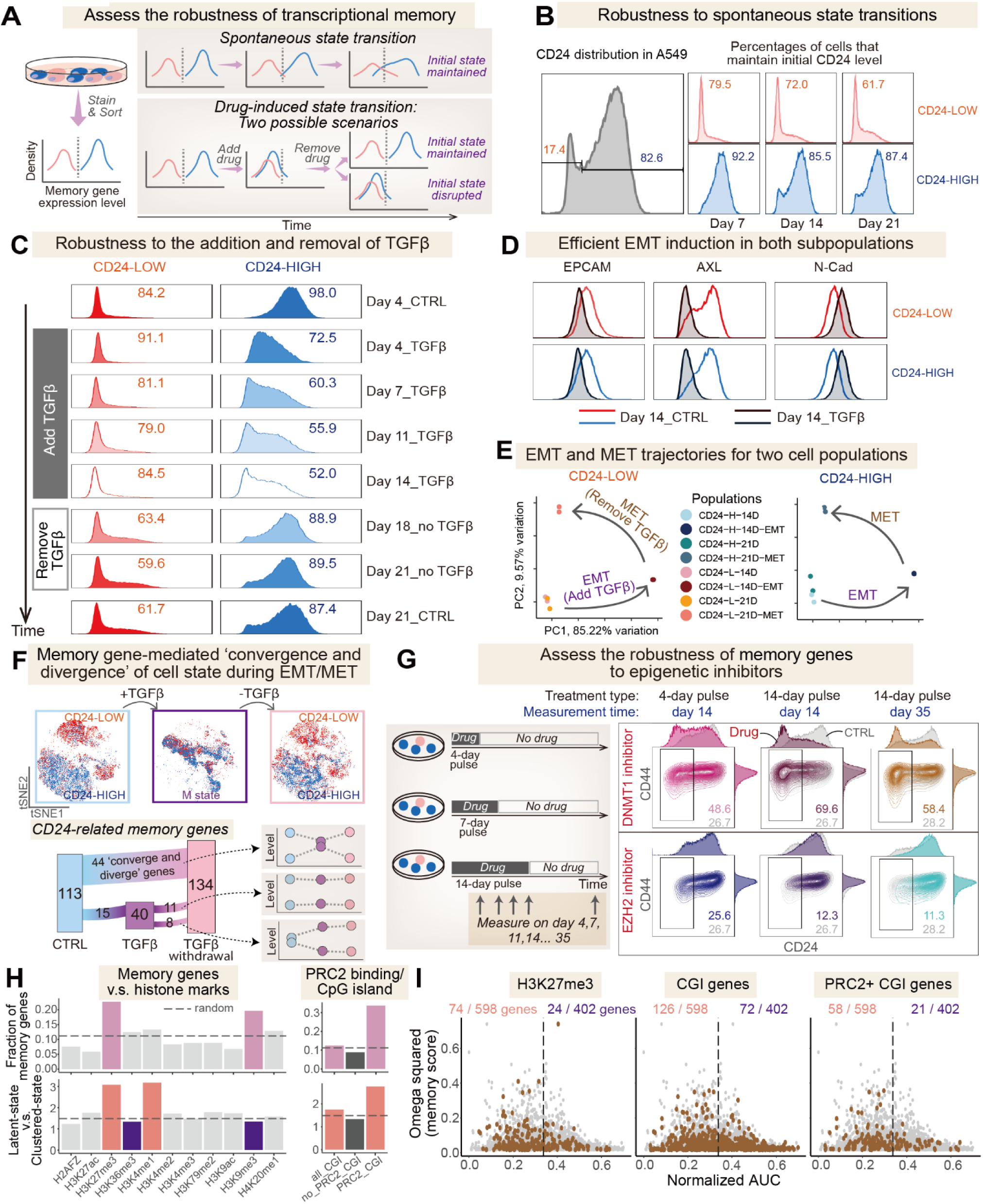
Epigenetic maintenance restores initial heritable states following transient plasticity. **A**, Experimental design for assessing the robustness of memory genes to perturbations. Using CD24 as an example, cells with top 20% CD24 level (blue) and bottom 20% CD24 level (red) are sorted and cultured separately. Subpopulations are cultured to observe spontaneous state transitions without treatment. For studying stimulus-induced state transitions, cells are treated with pulses of stimulus to test if initial variation between subpopulations can be maintained. **B**, Time-series tracing of spontaneous state transition of CD24-low (red) and CD24-high (blue) subpopulations. Most cells maintained their initial CD24 level for more than 21 days. **C**, CD24 distribution of CD24-low (red) and CD24-high (blue) subpopulations during TGF-β induced EMT for 14 days and subsequent MET for 7 days, measured by immunofluorescence staining and flow cytometry. CD24-low cells maintained their initial low CD24 level, while CD24-high cells showed significant down-regulation of CD24 during EMT and recovered their CD24 level after MET. **D**, Response of typical EMT markers during TGF-β treatment. CD24-low (red) and CD24-high (blue) cells showed up-regulation of N-Cad and down-regulation of EPCAM and AXL after TGF-β treatment for 14 days, measured by immunofluorescence staining and flow cytometry. **E**, PCA projection of global EMT/MET trajectory of CD24-low (left) and CD24-high (right) subpopulations, suggesting efficient and similar EMT/MET dynamics between subpopulations. EMT samples were collected at Day 14, and MET samples were collected at Day 21, along with paired control samples, all of which were measured by RNA-seq. **F**, Memory gene-mediated convergence and divergence of EMT trajectory for A549 cells. tSNE projection of multi-marker flow cytometry measurement representing the EMT trajectory of A549 cells. CD24-low cells (red) and CD24-high cells (blue) enter a similar “M” state (purple) after TGF-β treatment for 14 days and return to their initial state after TGF-β withdrawal for 7 days. The Sankey graph reflects the dynamics of *CD24*-linked memory genes in the EMT/MET process identified from RNA-seq in (**e**). Among these genes, a relatively large fraction (44 genes) displayed a convergence and divergence behavior analogous to *CD24*. **G**, Epigenetic perturbation experiments on the *CD24* memory state in A549 cells. EZH2 inhibitor (upper) or DNMT1 inhibitor (lower) was added to A549 cell cultures as a pulse of 4, 7, and 14 days, and CD24/CD44 levels were measured using time-series flow cytometry. **H**, Fraction of memory genes (top) and the ratio between latent-state and clustered-state genes (bottom) in A549 genes with indicated histone modification patterns and PRC2 binding/CGI-enriched patterns. Epigenetic patterns with specific enrichments were highlighted (light purple: patterns enriched for memory genes; dark purple: patterns enriched for clustered-state genes; orange: patterns enriched for latent-state genes). **I**, Association modes of memory genes enriched for indicated epigenetic patterns, including H3K27me3-enriched memory genes (n=98), CGI-enriched memory genes (n=198), and PRC2+/CGI-enriched memory genes (n=78).

We sorted A549 cells by CD24 surface expression into high (top 20%) and low (bottom 20%) populations, then cultured them with or without a 14-day TGF-β pulse. We measured EMT marker levels through flow cytometry and profiled transcriptomes by bulk RNA-seq (**Figures S7A-D**; **Table S10**; **Methods**). In untreated controls, we confirmed the memory of CD24 and other latent-state genes correlated with it. More specifically, we observed that, in the absence of TGF-β, most cells retained their initial CD24 levels with infrequent transitions (**Figure 6B**). Moreover, many genes differentially expressed between the CD24-high and CD24-low groups (**Table S10**) were also latent-state genes and enriched in EMT-related gene set (**Figures S7D-E**).

Strikingly, despite starting with different expression states of memory genes, both CD24-high and CD24-low subpopulations converged to a similar CD24-low “M” (mesenchymal) state upon TGF-β exposure, and then reverted to their original CD24 levels once TGF-β was removed (**Figure 6C**). Flow cytometry analyses showed the expected increase in N-Cad and decrease in EPCAM for both groups after EMT induction (**Figure 6D**), while bulk RNA-seq revealed comparable EMT trajectories in principal component space and similar changes in EMT-related genes during the whole EMT/MET process (**Figures 6E**, **S7F**).

Thus, after a transient but efficient TGF-β pulse, these two initially distinct “E” (epithelial) subpopulations converge to an “M” state before reverting to their original states (**Figures 6F**, **S7C**), consistent with the hypothesis that the memory of memory genes is robust to cellular perturbations. Intriguingly, a closer look (**Table S10**) indicated that many latent-state genes, including *CD24*, experience both convergence and divergence during the EMT/MET process (**Figure 6F** bottom, **Figures S8A-B**). Mechanistically, this observation aligns with the picture that the epigenetic states, rather than sheer expression levels, of memory genes are preserved, consistent with reports that DNA methylation patterns remain stable during TGF-β-induced EMT^64^. In other words, it appears that transient TGF-β stimulation alone does not erase the epigenetic memory of these genes. If that were the case, then perturbing the epigenetic states of the cell might more effectively disrupt their memory. To test this, we treated A549 cells with short-term pharmacological inhibitors targeting EZH2 (an H3K27me3 writer) or DNMT1 (a DNA methylation writer). Both treatments produced lasting effects: inhibiting EZH2 upregulated CD24, while inhibiting DNMT1 led to its downregulation (**Figures 6G**, **S8C**), providing support the picture that epigenetic modifications underpin the memory of latent-state genes.

Based on these results, we hypothesized that the two types of memory genes may be subjected to divergent epigenetic regulation. To test this, we analyzed the enrichment of histone marks of memory genes found in A549 and found that they were often enriched for H3K27me3 and H3K9me3 (**Figure 6H** top). In particular, H3K27me3 was more enriched on latent-state genes (**Figure 6H** bottom). Furthermore, expressive genes with high CGI (CpG island) content interacting with the PRC2 complex (a H3K27me3 writer) were more prone to have high memory and associate with latent states (**Figure 6I**), consistent with a picture that crosstalk between H3K27me3 and DNA methylation could orchestrate memory states of latent-state genes. Intriguingly, we found that in mESCs, latent-state genes were enriched for distinct epigenetic marks (**Figures S8D-E**) that mirror the observation in A549 cells (**Figures 6H-I**).

Together, these results are consistent with a picture that the memory status of memory genes is robust to perturbations, maintained by epigenetic marks, and can be erased by inhibitors of epigenetic writers. Moreover, evidence suggests that a conserved epigenetic regulatory principle may underly the memory of the two types of memory genes across both human and mouse cells.

## Discussion

Growing evidence underscores the biological relevance of heritable cellular heterogeneity^8–13^, but standard single-cell transcriptome datasets lack temporal information, hindering our ability to fully characterize its mechanistic basis and functional implications. Our study overcomes this limitation by leveraging temporal information implicitly encoded in lineage tracing data. We systematically demonstrate how pervasive and conserved transcriptional memory shapes heritable cell states, revealing insights into functional cellular heterogeneity and the roles these states play in cancer and developmental systems.

By “stitching” together pieces of landscape defined by clones of cells to reconstruct a full cell state landscape (as motivated by potential landscape theory^65,66^, see Supplemental Note 3), CORAL offers distinct advantages for profiling cell states (see Supplemental Notes 1 and 2 for detailed comparisons with experimental and computational counterparts). First, unlike conventional approaches that identify cell states using single-cell transcriptome data^19,26,20^, our approach does not rely on prior biological knowledge or data pre-processing. Moreover, while conventional SNN clustering requires researchers to select resolution parameters that often lead to over- or under-clustering without biological justification^20^, CORAL’s clone-centric approach enables arbitrarily fine resolution where each clone represents a potential cell state defined by heritable transcriptional programs. As such, the distinctions between the states defined by CORAL are more biologically pronounced than conventional approach (as exemplified by **Figure 1F**). Second, by design CORAL readily distinguishes stable, heritable states from transient states^19^, enabling mechanistic and functional dissections of meaningful variations within a cell population. Third, CORAL overcomes a major limitation in lineage tracing by extracting both cell state information and state transitions from single-timepoint clonal data, reducing the need for challenging dynamic sampling *in vivo*^62,67^.

We have successfully applied CORAL to identify biologically meaningful states in both cancer and stem cells. For cancer cells, we revealed *SOX2*-regulated heritable state in lung cancer cells, and uncovered a *ARF5*-mediated hidden regulatory axis of drug resistance in melanoma cells. For stem cells, we identified the *Rian*+ state of mESCs, which may delineate the clonal variation in developmental potential^53^, and showcased the robustness of the approach through the precise classification of HSC clones into distinct priming states without needing any prior information or pre-processing steps (as compared to existing work^43^). More importantly, we clearly demonstrated that for latent heritable states, subtle transcriptomic variations between latent states can have profound biological outcomes, underscoring the necessity for high-resolution reconstruction of heritable cell states for fully understanding functional heterogeneity.

Mechanistically, our clone-oriented analysis enables the dissection of the molecular basis of heritable states. The systematic, unbiased profiling of gene expression heritability led to a major discovery that transcriptional memory is pervasive and is conserved across conditions, cell types, and even species. We systematically linked such widespread transcriptional memory to heritable states through gene-specific ANOVA analysis along the cell state hierarchy, revealing both clustered-state genes and latent-state genes. Notably, the conservation of both memory and AUC across conditions and cell types underscores the role of gene-intrinsic mechanisms in conferring transcriptional memory^68^. Furthermore, results from phenotype mapping experiments illustrated the unexpected predictive power of latent-state genes, highlighting the role of these (typically) hidden memory genes that stratify critical cell states. These findings markedly advance our current understanding of transcriptional memory, suggesting the pervasive role it may play in diverse cell types. More generally, our framework provides the means to test a fundamental hypothesis: that pre-existing, heritable variations in gene expression are a key driver of differential clonal responses in diverse biological systems^6,69^.

Intriguingly, our investigation into the epigenetic basis of these two classes of memory genes provides compelling justification for the conceptual utility of the “epigenetic landscape”^27,70^ in depicting cell state dynamics. Because the two memory gene classes appeared to be regulated by divergent epigenetic mechanisms, with latent-state genes likely being orchestrated by H3K27me3 and DNA methylation, the two types of heritable states defined by these genes are thus epigenetically associated. These findings also open avenues for in-depth study of the roles of chromatin binding proteins in shaping transcriptional memory and heritable cellular variations. Overall, our integrated workflow provides a powerful toolkit to systematically dissect cellular heterogeneity, enabling robust investigations into its pervasiveness, mechanisms, and functions across biology.

## Methods

### Cell culture

A549 cells (BNCC, Bena culture collection) were maintained in RPMI 1640 medium (Gibco, #C11875500CP) supplemented with 10% fetal bovine serum (FBS) (Excell Bio, #FSP50) and 1% Pen Strep Glutamine (Gibco, #10378016). HEK293T cells were maintained in Dulbecco’s Modified Eagle Medium (Gibco, #C11995500CP) supplemented with 10% fetal bovine serum (FBS) (Excell Bio, #FSP50) and 1% Pen Strep Glutamine (Gibco, #10378016). All cell lines were incubated at 37°C with 5% CO2-air atmosphere with constant humidity and routinely tested for mycoplasma. We passaged cells with 0.05% trypsin-EDTA (Gibco, # 25300062). Passage numbers were kept and cultures past a total of 25 passages were discarded.

### Plasmid construction

If not specified, plasmids used in this research were constructed using Gibson assembly methods. Oligonucleotides were synthesized by GENEWIZ. Backbones were linearized using restriction endonucleases from NEB. PCR fragments were produced using PrimeSTAR Max DNA Polymerase (TAKARA). All plasmids were confirmed by Sanger Sequencing (RUIBO).

### Barcode lentivirus library construction

For the barcode library, the vector was constructed based on pHR lentivirus backbone by Gibson assembly as described above. Specifically, human *EF1a* promoter, *EGFP* and *BGH* polyadenylation sequence are cloned into the digested vector and XmaI and PacI enzyme sites are added in 3’UTR of *EGFP*. Then we digested the vector by XmaI (NEB) and PacI (NEB) overnight and gel purified the linearized backbone. An 89 bp random barcode oligonucleotide with flanking sequences and a 24 bp reverse primer are ordered from GENEWIZ. Then we performed an extension reaction to generate dsDNA barcode for assembly: 5 μL KAPA hotstart HIFI Buffer (high-fidelity), 0.75 μL KAPA dNTP Mix, 0.75 μL 100 μM barcode, 1.5μL 100 μM reverse primer, 0.5 μL KAPA hotstart HIFI DNA Polymerase and 16.5 μL water. The reaction is incubated as below: 98 °C for 2 min, 10× (65 °C for 30 s, 72 °C for 10 s), 72 °C for 2 min, hold at 4 °C. dsDNA barcode was purified by columns (QIAGEN HiPure Gel Pure Micro Kit, Magen, D2120). We used Gibson assembly to combine the linearized backbone and barcode in over 1:20 ratio, purified the Gibson product by columns as before and transformed the reaction into Endura™ ElectroCompetent Cells. Transformants were recovered in Recovery Medium for 1 h in 30 °C before plating several dilutions and seeding culture in 50 mL LB with 100 µg/ml carbenicillin in 30 °C. Barcode insertion was verified by colony PCR reaction and Sanger sequencing. Plasmids from separated cultures are pooled before lentivirus production.

### Lentivirus packaging

For lentivirus transfection experiments, we grew 293T cells to about 85% confluency in 6 well plates. We combined 200 μL Opti-MEM (Gibco, 31985070) with 2.66 μg PCMV plasmid, 0.34 μg PMD plasmid, 3 μg transfer Plasmid and PEI in one tube such that the ratio of μg DNA:μg PEI is 1:4. We vortexed the tube incubated the mixture at room temperature for 30 min and pipetted the solution dropwise into the plate of 293T. After incubation for 6∼8 h, we removed the media and added 3 mL of media and harvest media containing virus at 48h post transfection. We centrifuged the viral supernatant at 500 g for 5 min and filtered through a 0.45 μm PES filter before storing them at −80°C.

### Lineage tracing experiment of A549 cells *in vitro*

Cells were seeded in a well of six-well plate and transfected with the barcode lentivirus pool with <0.1 MOI. Cells were incubated with virus overnight and grew for 2∼3 days to start expressing the barcodes. For A549 cells, ∼10000 barcoded cells were sorted and allowed to expanded for 3∼4 doublings. Then we plate ∼500 cells on a well of 96-well plate from the barcoded cell pool and expand them for ∼7 days before performing single-cell RNA-seq experiment. The rest of the cells in the barcoded cell pool were expanded parallelly for 10 days and cryopreserved in 4 vials with 10^7^ cells each for further phenotype mapping experiment.

### Lineage tracing experiment of A549 cells *in vivo*

Our animal experiments were performed in accordance with institutional and national guidelines. The protocols (AAIS-LinYH-1) have been approved by the Peking University IACUC. A549 Cells were seeded in a well of six-well plate and transfected with the barcode lentivirus pool with ∼0.1 MOI. After incubation with virus overnight, ∼10^5^ transfected cells (with ∼10^4^ barcoded cells) were mixed with 5×10^6^ WT cells and injected into a 4–5-week-old BALB/c nude mouse. Tumor growth was monitored twice a week till the volume of tumor achieved 1000 mm³. Then, the mouse was euthanized and the tumor tissue was extracted for further analysis.

### Single-cell RNA-seq

For lineage tracing experiment *in vitro*, we trypsinized cells, resuspended them in DPBS and followed the protocols of the Chromium Next GEM Single Cell 3ʹ Reagent Kits v3.1 (Dual Index) (10X Genomics). For lineage tracing experiment *in vivo*, we used the Besttopcell mammalian cell tissues disassociation kit (BA3303) to get single-cell suspension of tumors, used MoFlo XDP (BeckMan Coulter) to sort GFP+ barcoded cancer cells and then performed the same protocols of Chromium. The cell suspension was passed through a 70 µm nylon cell strainer (ThermoFisher Scientific, 22-363-548) to remove clumps and debris, centrifuge the cells at 300 g for 5 min at 4 °C, discard the supernatant and the cells were re-suspended in PBS (Bioss, C7033) containing 5% FBS (Solarbio, s9020). Then cells were counted at count star. We observed that cell contained red blood cells in the count star instrument, next, we used red blood cell lysis buffer to remove red blood cell. Following a 15 min incubation at 4 °C, samples were centrifuged at 450 g for 5 min at 4 °C, discard the supernatant and wash twice times using PBS containing 5% FBS, Cells were then re-suspended in PBS containing 5%FBS. Next, cells were counted at count star and adjust the cell suspension concentration (600-1200 live cells per microliter). Then we followed similar protocols of the Chromium Next GEM Single Cell 3ʹ Reagent Kits v3.1 (Dual Index) (10X Genomics) to finish single-cell RNA-seq.

### Computational analysis of single-cell RNA-seq

Raw reads of single-cell RNA-seq were demultiplexed and mapped to the reference genome by 10X Genomics Cell Ranger pipeline using default parameters. The Cell Ranger software was obtained from 10X Genomics website. Alignment, filtering, barcode counting and UMI counting were performed with cell ranger count module to generate feature-barcode matrix.

We performed the downstream single-cell RNA-seq analysis on Seurat V5^71^. For lineage tracing experiment *in vitro*, genes within less than 3 cells and cells with less than 200 genes or >5% mitochondrial gene fraction were first removed. Then we performed data dimension reductions including PCA and UMAP and SNN clustering using the default settings of Seurat. The cell cycle score was calculated by CellCycleScoring function of Seurat and the cell cycle effect was removed by regressing out the difference between the G2M and S phase scores in the ScaleData function get the cell cycle-regressed UMAP visualization.

For the lineage tracing experiment *in vivo*, genes within less than 3 cells and cells with less than 200 genes or >5% mitochondrial gene fraction were first removed. Then we performed data dimension reductions including PCA and UMAP and SNN clustering using the default settings of Seurat. The cell cluster with immune-relevant gene expression including *ZNF90* and *CXCL14* were firstly removed from the sample. Then we performed data dimension reductions including PCA and UMAP to the preserved cells for further CORAL analysis.

See **Table S11** for a summary of all lineage-tracing single-cell RNA-seq datasets used in this study.

### Cellular barcode recall from single-cell RNA-seq

For lineage tracing experiment *in vitro*, cellular barcodes were extracted directly from the bam files of unmapped reads generated from Cell Ranger using pycashier. For lineage tracing experiment *in vivo*, we used excess full length cDNA sample from single-cell RNA-seq experiment workflow to improve the recovery of cellular barcodes. Specifically, we used a side PCR reaction, targeting the 3’UTR region between GFP and cellular barcodes. ∼100 ng cDNA, side PCR primers (10 μM) and KAPA HiFi HotStart ReadyMix were mixed and we used the following protocols: 98 °C for 30 seconds, 24 cycles of 98 °C for 10 seconds and 65 °C for two minutes, followed by a final step at 65 °C for five minutes. Then the product was purified by 1.2x VAHTS® DNA Clean Beads and the NGS of the amplified barcodes was conducted by Annoroad. Then the connection between 10X cellular barcodes and clone barcodes were counted simply from the NGS data.

To improve the barcode recovery accuracy from lineage tracing data, we performed a filtering step to assign cells to unique clone barcodes. For the lineage tracing experiment *in vitro*, we only preserved clone barcodes with >300 raw counts. Each cell was assigned to its dominant clone barcode, the count of which is twice as large as the second largest barcode in the cell. If there is no significant clone barcode, the cell would be assigned to observed barcode that is most abundant in the whole dataset and observed in the cell.

### Simulation of cell state transition with multi-stage cell cycle model

The simulation of cell state transition and CORAL reconstruction was implemented by MATLAB. We simulated the distribution of cell populations generated from a specific initial cell. The proliferation of cells is modelled as Erlang process, which assumes that cells need to go through *K* cell cycle phases with similar length following Poisson distribution^72^.

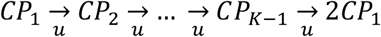

Thus, for the initial cell *I*_*i*,1_ with heritable state *i*, the number of its progeny can be simulated by Gillespie algorithm according to the following

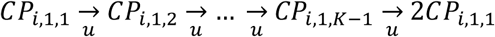

When the potential transition between cell state *i* and cell state *j* happens, we have:

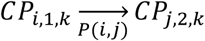

Recursively, the progeny of the new generated sub-initial cell *I*_*i,l*_ (*l*=current sub-clones number) can be simulated as Gillespie algorithm as well.

In short, the output of simulation is hence the number of *CP*_*i,l,k,t*_ (*i* ∈ {*cell state*}, *l* ∈ {*sub* − *lineages*}, *k* ∈ {1,2, … *K*}, *t* ∈ [0, *T*]. Then the number of cells with specific state i at time *t* can be calculated as:

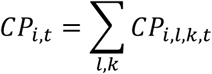

To simplify the simulation process, we used the average cell cycle length as time unit in the following simulation analysis.

### CORAL analysis of simulated clonal data containing hidden cell states

The simulation for evaluating CORAL’s performance was implemented by MATLAB. Specifically, in the toy model of cell state transition, we assume that each cell pair from specific cell state combinations has its characteristic cell-cell distance ***D***_*ij*_. We noted that the cell-cell distance is not necessarily proportional to the cell state transition matrix ***P***_*ij*_. As we could get the distribution of different cell state of a clone from simulation mentioned before, now the distance between clone ***m*** with m cells and clone ***n*** with n cells could be calculated as:

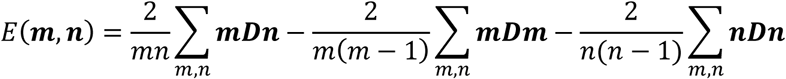

which is the *U* statistic, an unbiased estimation of Energy distance^28^. We adjusted the negative E to zero so that it can be treated as distance matrix.

For visualization of clones, we performed classical MDS^29^ to the distance matrix. For identifying heritable cell states, we performed hierarchical clustering with Ward methods^30^ to obtain cell state divisions of clones. The performance of CORAL simulation was evaluated by the proportion estimation accuracy and the initial cell state assignment accuracy.

### Simulation of stochastic cell state transition and imperfect clone sampling

For the demonstration of hidden cell state detection task, we set three cell states: state A, B and C. The transcriptome distance between A and B was 0 and the transition rate between state A and state B was much slower than the transition between state B and state C (**Table S1**). The time-series data of cell state distribution was generated by MATLAB from 500 clones, which were assigned to random initial states sampled from steady-state distribution.

For the parameter scanning of cell state transition rate, the cell states were set as before and we scanned the cell state transition timescale parameter space. We generated distribution of cell states from 500 clones on each parameter combination for five times to estimate the performance of CORAL.

For the parameter scanning of imperfect clone sampling, the parameters were set as the default hidden cell state detection task, and the clones and cells per clone were down sampled according to the parameter settings.

### CORAL workflow for single-cell RNA-seq data in A549 cells

We first filtered out cells without barcodes and clones with <3 cells. The energy distance between clones were calculated as described before:

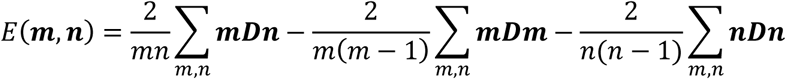

where *D*(*i*, *j*) is the Euclidean distance between cells.

The CORAL tree was reconstructed using hierarchical clustering of “Ward.D” method with all clones remained. The MDS visualization was then generated using the classical MDS algorithm. See Supplemental Note 3 for more details.

For lineage tracing experiment *in vitro*, clones were clustered into 2, 6 and 24 heritable states to evaluate CORAL’s performance in different cell state granularity. Then the cell cycle phase distribution of each heritable state and the DE genes of each heritable state were calculated by the CellCycleScoring and FindMarkers functions of Seurat. For lineage tracing experiment *in vitro*, only G1 phase cells were preserved for DE gene detection to minimize the cell cycle effects. For lineage tracing experiment *in vivo*, clones were clustered into 3 heritable states.

The variance explained by heritable cell states under various cell state resolution was measured by the eta squared and omega squared:

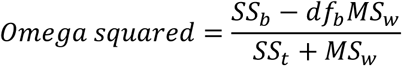

And we noted that in the Euclidean space, the *SS_w_* and *SS*_*t*_ can be written in the form 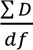 so the omega squared can be calculated from the cell-cell distance matrix.

### Memory gene identification, association mode analysis, and related analyses

The “data” slot of the Seurat object was used for gene-level statistical analysis. For the memory gene analysis, we first selected genes with >0.05 mean expression. Then the omega squared of ANOVA test was calculated as below:

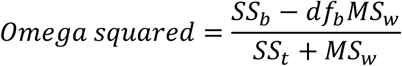

where *SS*_*b*_ is sum of gene variance between clones and *SS*_*t*_ is the sum of gene variance in total. *df*_*b*_ equals the clone number and *MS*_*w*_ is the mean square between clones. We used randomly shuffled “fake” clone identity to calculate the fake omega squared values over five repetitions. The final cutoff for omega squared was set as the third quartile (Q3) plus 1.5 times the interquartile range (IQR) of the fake omega squared values. Genes with omega squared values above this cutoff were selected as memory genes.

For the association mode analysis, we performed another ANOVA analysis on the memory genes using the heritable state definition (i.e., CORAL state) instead of clone identities as groups. We calculated the approximate AUC shape as below:

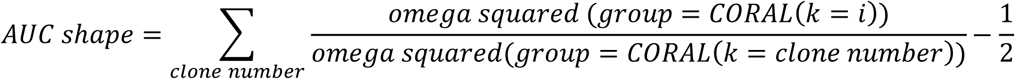

The global AUC shape was calculated similarly. Genes with larger AUC shape than the global AUC shape were designated as clustered-state genes while genes with smaller AUC shape than global AUC shape were designated as latent-state genes.

The pseudobulk expression matrix for co-expression analysis of memory genes were generated by the AggregateExpression function in Seurat and then normalized. The gene list of ENCODE Histone modifications properties was extracted using the “A549” relevant terms of “ENCODE Histone Modifications 2015”^73^ dataset from the R package enrichR while the gene list of CGI-enriched gene was extracted from previous publications^74^.

### Comparation with Memory-seq for identifying memory genes

We compared our approach for memory gene identification with Memory-seq^9^. We adapted the method from the Memory-seq analysis of bulk RNA-seq data previously reported^9^, which used average expression level and expression variance at clone level to extract genes with higher variance than noise. In our comparison, we used the AggregateExpression function in Seurat to generate pseudo-bulk expression matrix of lineage tracing dataset to perform Memory-seq analysis.

More specifically, the Memory-seq workflow fits the Poisson regression model between the mean expression and CV, and genes with high residual are treated as high memory genes. We noted that the analytical Memory-seq analysis assumes that the on rates of memory genes are rare, and thus would in principle lead to a systematic underestimation of memory genes with relatively high on rates, such as *CD24*.

### Identification of cell cycle genes and ribosomal protein genes

Cell cycle genes were identified by the cc.gene function of Seurat. The ribosomal protein gene list was downloaded from the Ribosomal Protein Gene Database (http://ribosome.med.miyazaki-u.ac.jp/rpg.cgi?mode=orglist&org=Homo%20sapiens).

### CORAL analysis on WM989 dataset

The public WM989 lineage tracing dataset^11^ were downloaded from the https://drive.google.com/drive/folders/1-C78090Z43w5kGb1ZW8pXgysjha35jlU?usp=sharing in the form of 10×1_Filterd_BatchCor_unnorm_sctrans.rds. We used pre-processing Seurat object, SNN clustering result, the assigned clone barcode identity and initial state identity (*EGFR*/*NGFR*+ or not) of original publication^11^.

For the CORAL analysis, cells without clone identity or in small clones (<2 cells) were filtered out before calculating the clone-clone distance matrix. 321 clones were preserved for the reconstruction of heritable states and MDS visualization. 9305 memory genes were identified from 9483 expressive genes. Top 500 memory genes were preserved for co-expression analysis.

For the analysis of crosstalk between ARF5 and other memory genes. Clones were divided as “*ARF5*-high” and “*ARF5*-low” clones by the median expression level of *ARF5* and the *EGFR*/*AXL*/*NT5E* expression level were compared by Wilcoxon rank sum test in R.

### CORAL analysis on mESC dataset

The public mESC lineage tracing dataset^51^ was available at the Gene Expression Omnibus (GEO) under accession number GSE226169, and we used the assigned clone identity of original publication. For the CORAL analysis, the largest 5 clones with extremely large clone sizes after 48h culture were removed and the clone with at least 2 cells were remained. 247 clones were preserved for the reconstruction of CORAL and MDS visualization. 1044 memory genes were then selected from 8765 expressive genes. For the epigenetic regulation pattern analysis of memory genes, the histone modification properties of mESCs were extracted using the “ES-Bruce4” relevant terms of “ENCODE Histone Modifications 2015” dataset from the R package enrichR while the gene list of CGI-enriched gene was extracted from previous publications as well.

### CORAL analysis on HSC lineage tracing dataset

The public HSC lineage tracing dataset^43^ was available at figshare (DOI: 10.6084/m9.figshare.25822948). We used the D7/female donor subset of the HSC *in vitro* culture lineage tracing dataset and used the assigned clone identity and cell type identity of original publication. The UMAP embeddings were generated as described as original publication, which used all samples and removed the effect of cell cycle and ribosomal genes.

For the CORAL analysis, cells without clone identity or in small clones (<2 cells) were filtered out before calculating the clone-clone distance matrix. The preserved 841 HSC clones at Day 7 were divided into 4 types using hierarchical clustering with Ward method. DE genes of HSC heritable state under specific cell type were extracted using FindAllMarkers function of Seurat with default parameters and a 0.05 cutoff of adjusted p-value.

For the primed and origin markers detection in specific cell types, we compared cells between different heritable states but the same cell type. We excluded the monocyte, neutrophil, basophil and ery cells as their cell numbers were too few. Meanwhile, heritable state with less than 200 cells in specific cell type was excluded from the analysis to reduce the noise in the data. The DE genes were calculated by FindAllMarkers function in Seurat with default parameters and a 0.05 cutoff of adjusted p-value.

For memory gene identification, 1388 genes were selected from 9142 expressive genes as memory genes, which were used for further association mode analysis.

### Circular visualization of memory genes across datasets

To visualize the landscape of memory genes, a circular plot was generated using the R package circlize. Memory genes were first subjected to functional annotation. Gene sets from MSigDB^75^ (including Hallmark, KEGG, Reactome, and Gene Ontology categories: Biological Process, Cellular Component, and Molecular Function) were fetched using the msigdbr package. A keyword-based mapping strategy was employed, where each memory gene was associated with pathways it belonged to. A predefined, prioritized list of keywords (e.g., “HALLMARK_HYPOXIA” for Hypoxia, “GOBP_CELL_CYCLE” for Cell Cycle) was used to assign genes to a primary functional category; more specific keywords and curated pathways (e.g., Hallmarks) took precedence over broader terms. Ribosomal protein genes were specifically annotated based on established gene name patterns (e.g., *RPL*, *RPS*). Genes not matching any defined keywords were categorized as “Other”. For comparative analysis, the memory status of memory genes identified in A549 *in vitro* was assessed in four other datasets. Mouse gene names from mESC and HSC datasets were mapped to human orthologs using R package homologene, with simple capitalization as a fallback for unmapped genes.

### Migration assay

Before performing phenotype mapping experiment, the cryopreserved parental barcoded cells were thawed and cultured for at least 48 h to recover their viability. Specifically, migration assay in the phenotype mapping experiment, 2.5×10^6^ parental barcoded cells were washed twice by DPBS and added to the insert of each well of Costar Transwell plates (Corning, #3428) in 0.6 mL serum-free complete medium, with 1mL complete medium in the receiver well. Cells were then incubated for 18 h at 37 °C. The top and the bottom of the transwell inserts were washed twice with DPBS and Trypsin was added to the insert (0.6 mL) and the receiver well (1 mL). Complete medium was added to inactivate the Trypsin and cell suspension from inserts and receiver wells were collected. Cells from 4 wells were combined for further analysis.

### 3D culture assay

For 3D culture assay in the functional mapping experiment, 2.5×10^6^ parental barcoded cells were plated for 3D culture in complete medium supplemented with 1.5% Matrigel (Corning, #354234) in each 10cm ultra-low attachment dish (Labselect, #12331). Media was replenished every four days by aspirating 50% of the media and replacing with fresh complete media supplemented with 1.5% Matrigel. Spheroids grown for 7 days were collected by Accutase (ThermoFisher, #A1110501) treatment for 15min. Cells from 4 wells were combined for further analysis.

### Drug resistance assay

For drug resistance assay in the functional mapping experiment, 2.5×10^6^ parental barcoded cells were plated in each 10 cm dish and cultured for 24h. Then gemcitabine (Cayman, 11690) was added to the culture in 10 μM final concentration. The survived cells were collected after 72h. Cells from 4 well were combined for further analysis.

### Immunofluorescence imaging

A549 cells were plated on 24-well glass-bottom plates for 4 days before staining. Cells were fixed in 4% paraformaldehyde for 30 min and then washed 5 times with PBS after fixation. Cells were then blocked in 1% BSA in PBS for 1 h and incubated with primary antibodies (purified anti-human CD24 Antibody, 1:1000, ML5, BioLegend) in 1% BSA at 4 °C overnight. Cells were washed 5 times and then incubated with secondary antibodies (donkey anti mouse IgG (H+L) Highly Cross-Adsorbed Secondary Antibody, Alexa Fluor Plus 647, 1:1000, Invitrogen) and DAPI (75 μg/mL, Beyotime) in 1% BSA at room temperature for 1 h. Cells were then washed 5 times with PBS and imaged under a Leica TCS SP8 confocal laser scanning microscope. The samples were protected from light for every step.

### Barcode quantification from gDNA

Genomic DNA of barcoded cell population was extracted using Quick-DNA™ Miniprep Plus Kit (Zymo). Then the barcode sequences were amplified using primers containing Illumina adapters and unique sample indices: 25 μL KAPA hotstart ReadyMix, 0.75 μL forward primer (10 μM), 0.75μL reverse primer (10 μM), 500ng gRNA and water up to 50 μL. The reaction is incubated as below: 95 °C for 2 min, 25× (98 °C for 20 s, 65 °C for 15 s, 72 °C for 15 s), 72 °C for 2 min, hold at 4 °C. We scaled up the number of reactions so that all gRNA can be amplified. The PCR products were then pooled and purified using HiPure PCR Pure Mini Kit (Magen, D2121) and the purified product was run on the gel before gel purification to remove primer dimers. Gel purification was conducted using HiPure Gel Pure Micro Kit (Magen, D2110). The NGS was conducted by Annoroad.

### Computational analysis of phenotype mapping experiment

The barcode count of each phenotype mapping experiment was estimated by pycashier^76^. The barcode counts were first normalized to the total counts of each sample and the relative abundances (functional score) were calculated by comparation to the normalized counts of control group. Clones with >250 raw counts (3147 clones in total) in the NGS data of control group were preserved for further analysis. For the LASSO feature selection, the largest 200 clones in the dataset of *in vitro* A549 cells were mapped with the relative abundances in the phenotype mapping experiment. The LASSO algorithm was conducted by the R package glmnet. Specifically, we ran the linear regression (family= “guassian” and set alpha=0.3. We ran cv.glmnet for 100 times to average the error curves in the cross validation to find the optimized lamda. Then we used glmnet and coef function to get the key features selected by LASSO.

### Computational analysis of lineage tracing dataset of lung cancer metastasis

The lineage tracing dataset of lung cancer metastasis^62^ was generated from GSE161363 and the data of the “m5k” mouse was used for further analysis. Cells sampled from left lung were preserved for classical single-cell analysis and CORAL analysis, and the static barcode “iniBC” was used to perform CORAL analysis. Genes within less than 3 cells and cells with less than 200 genes or >5% mitochondrial gene fraction were first removed. The cell cycle score was calculated by CellCycleScoring function of Seurat and the cell cycle effect was removed by regressing out the difference between the G2M and S phase scores in the ScaleData function get the cell cycle-regressed UMAP visualization. Then we applied CORAL workflow and clustered clones into 6 heritable states. The migration rate, or the “TreeMetRate” was estimated as described in original research and clones only detected in the left lung sample were assigned as non-migrative clones.

### Flow cytometry analysis and sorting

Cells were lifted off using Accutase (ThermoFisher, #A1110501), washed with Cell Staining Buffer (CSB, BioLegend, #420201) and resuspended in ice-cold CSB. For cell surface marker staining, the premixed antibodies were added according to the manufacturer’s instructions and incubated in the dark on ice for 30 mins. For EMT/MET tracing experiment of CD24 subpopulations, we used a mixture of CD24-BV421 (1:100, ML5, BioLegend), EPCAM-BV650 (1:100, 9C4, BioLegend), CD44-BV785 (1:100, IM7, BioLegend), AXL-PE (1:100, DS7HAXL, eBioscience) and N-CAD-PE-CY7 (1:100, 8C11, eBioscience). For the time-series epigenetic perturbation experiment, we used a mixture of CD24-PE (1:100, ML5, BioLegend) and CD44-APC (1:100, IM7, BioLegend).

Cells were then washed in CSB and resuspended in 300 μL CSB. For the EMT/MET tracing experiment of CD24 subpopulations, Sytox Green (Invitrogen™, S34860) were added to the CSB with 30 nM final concentration to exclude dead cells. For flow cytometry analysis, we used BD Fortessa SORP instrument for time-series epigenetic perturbation experiment and used Symphony A5SE instrument for complex EMT panels in EMT/MET tracing experiment. For FACS experiment, we followed the same protocol and sorted cells in BD Aria Fusion instrument. Sorted cells were centrifuged and resuspended as needed. The flow cytometry data is analyzed by FlowJo V10. The t-SNE projection of flow cytometry data is generated with all EMT markers under default parameters of FlowJo V10.

### EMT induction and epigenetic perturbation experiment

WT or sorted cells were seeded in 6 well plates at 300,000 cells/well. For FACS experiments, drugs were added 24 hours after seeding. During the drug treatment, we replaced media containing drugs every 2 days and passaged the cells as needed. The final TGFβ (Novoprotein, CA59) concentration for EMT induction is 10 ng/mL. For the epigenetic perturbation experiment, the concentration of tazemetostat (Selleck, #S7128) is 10 μM while the concentration of GSK-3685032 (Selleck, #E1046) is 500 nM.

### Bulk RNA sequencing and analysis

Cell samples were collected and lysed by Trizol (Invitrogen, 15596018) and stored at −80 °C. RNA extraction, library construction and sequencing were conducted by Annoroad. For data analysis of RNA sequencing, we used Salmon^77^ for quantifications and Tximeta^78^ for importing the abundance matrix into R. The PCA projection of RNA-seq data was generated by the R package PCATools and the DE genes were calculated by DEseq2^79^. DE genes with <10e-3 p.adj and >0.15 log2FC were preserved for further visualization and analysis.

### GSEA analysis of memory genes and DE genes in EMT/MET process

The gene set enrichment analysis (GSEA)^80^ of memory genes DE genes between CD24-low and CD24-high populations and DE genes during EMT/MET process was performed by the R package ClusterProfiler^81^. We used the human HALLMARK gene sets available in the MSigDB database^75^ (https://www.gsea-msigdb.org/gsea/msigdb). The GSEA algorithm used enrichment score to estimate the degree of overrepresentation at the top or bottom of the ranked list of DEGs. The positive enrichment score indicates gene set enrichment at the top of the ranked list while a negative score indicates enrichment at the button of the ranked list.

## Data and materials availability

Original code and data objects will be deposited at figshare and can be temporally accessed from here: https://drive.google.com/drive/folders/1-cNiSKZFyVSs9Mndq87AcRXfaGweLesj?usp=sharing. The codes are also available in https://github.com/YongjieLin1998/CORAL. The relevant sequencing data reported in this work will be deposited to NCBI. All other data and materials are available from the corresponding author upon requests.

## Author Contributions

Yongjie L. designed the project, performed experiments and data analysis, and wrote the manuscript; X.C. performed experiments and data analysis; L.W. performed experiments and data analysis; Y.Z. provided help to data analysis; Yihan L. designed and supervised the project, and wrote the manuscript; all authors contributed to reviewing and editing of the manuscript.

## Acknowledgments

We thank members of the Lin Lab, especially Yuanzhuo Wang, Yadi Liu, and Baoqiang Chen, for helpful discussions regarding the experiments, analysis, and manuscript. We thank the Center for Quantitative Biology at Peking University and the flow cytometry core at the National Center for Protein Sciences at Peking University for equipment support of Flow Cytometry Facility, particularly Zeng Fan, Fei Wang, Xuefang Zhang, Jia Luo and Liying Du for the technical help. We thank the Quantitative Imaging facility at the Center for Quantitative Biology at Peking University for equipment support, particularly Xin Li for the technical help. We thank the High-Performance Computing Platform of the Center for Life Science for computational support. AI-based tools were used for grammar and style checking. Yihan L. acknowledges grants from National Natural Science Foundation of China (Grant # T2325002, 32088101, and T2321001), and National Key R&D Program of China (Grant # 2020YFA0906900 and 2018YFA0900703).

## Declaration of interests

The authors declare no competing interests.

## Supplemental Information

Supplemental Figures 1-8

Supplemental Notes 1-3

Supplemental Tables 1-11

## Supplemental Figures

**Figure S1:**
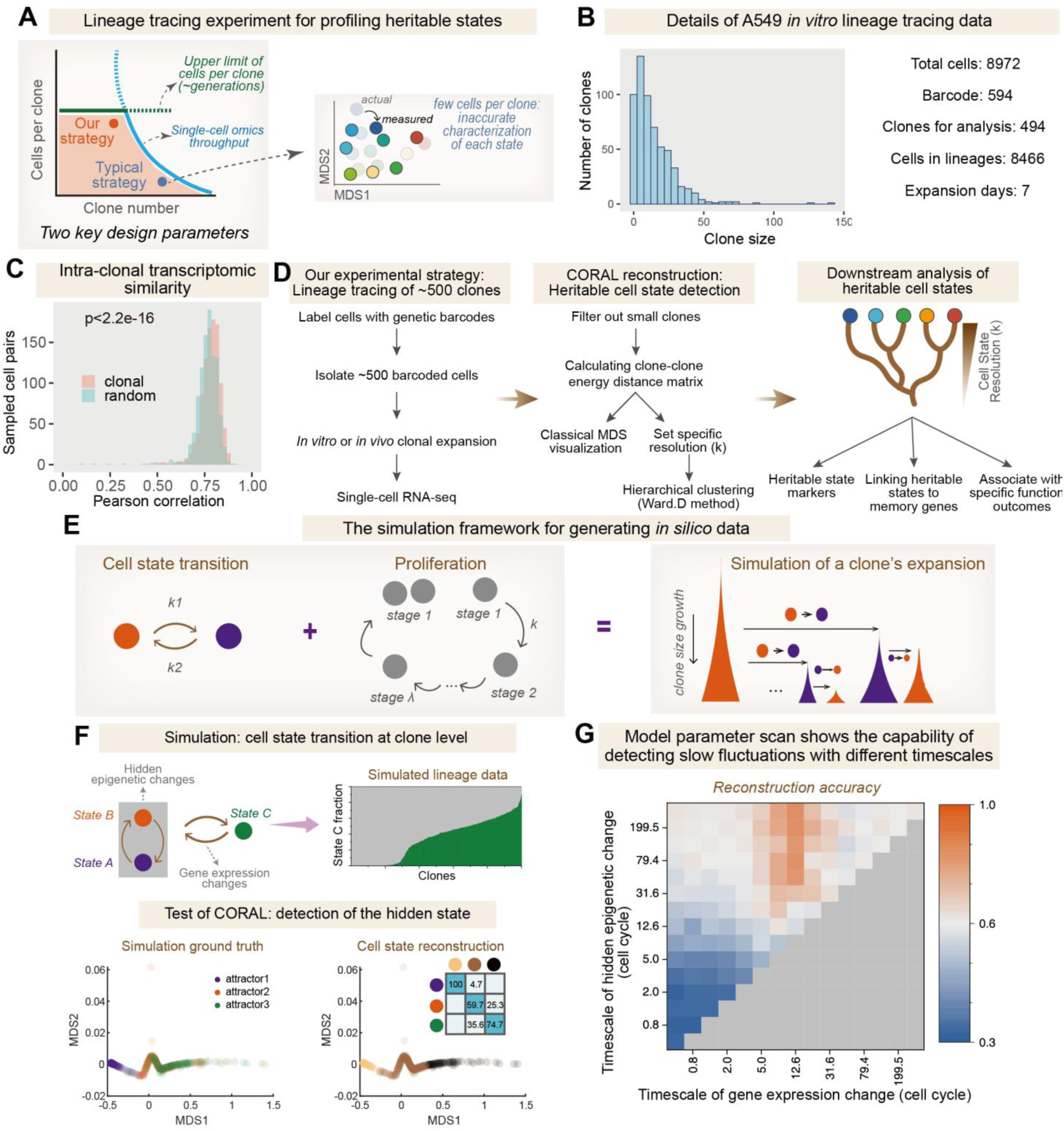
Experimental design, procedure, and in silico validation of CORAL. **A**, Schematic illustrating the design of lineage tracing experiment for profiling heritable cell states. The clone number and cells sampled per clone are limited by single-cell omics throughput, while the number of cells per clone is also constrained by the expansion generations of clones (left). Classical lineage tracing strategies often collect many clones with few cells per clone, leading to incorrect estimations of clones’ transcriptome distributions (right). In contrast, our experimental strategy reduces the clone number (to ∼500) to maximize the cells sampled per clone (i.e., dense clonal sampling). **B**, Distribution of clone sizes in the lineage tracing experiment of A549 cells *in vitro*. 494 clones with at least 3 cells (out of a total of 8466 cells from 594 total clones) were recovered after a 7-day expansion for further analysis. **C**, Pearson correlation of random cell pairs and cell pairs from the same clone. 1000 random and clonal cell pairs were sampled, and the p-value was calculated using a two-sided Wilcoxon test. **D**, Flowchart illustrating our experimental and computational workflow. Data are collected by performing single-cell RNA-seq on lineage-encoded cell populations starting from ∼500 progenitor cells. The clone-clone distance matrix is calculated to enable clone-level MDS visualization and heritable cell state reconstruction by CORAL. The resulting heritable states can be utilized for detecting memory genes, linking memory genes to cell states, and studying functional relevance. **E**, Framework for simulating cell state transitions, where cells in different states (red or blue) may stochastically transition to other states in a Poisson process (**Methods**). The proliferation process is modeled as an Erlang process, reflecting the multi-stage nature of the cell cycle^72^. Cell state transition and proliferation processes are simulated jointly, creating many proliferating “sub-clones” with specific cell state transition histories during a clone’s expansion process. **F**, Testing CORAL-based cell state reconstruction using simulated data. A three-state toy model with indistinguishable cell states (purple and orange) was employed to generate clone-level cell state distribution data, with only the proportion of the third state (green) observable (**Methods**). MDS embeddings of 1000 simulated clones successfully distinguished not only the visible state (green) but also the hidden states (purple and orange). CORAL’s clustering method accurately assigned most clones to their correct initial states, as shown in the confusion matrix. **G**, Parameter scanning of CORAL’s capability under different kinetics. A heatmap illustrating the accuracy of initial cell state assignment by CORAL under varying timescales in the cell state transition model presented in (**F**).

**Figure S2:**
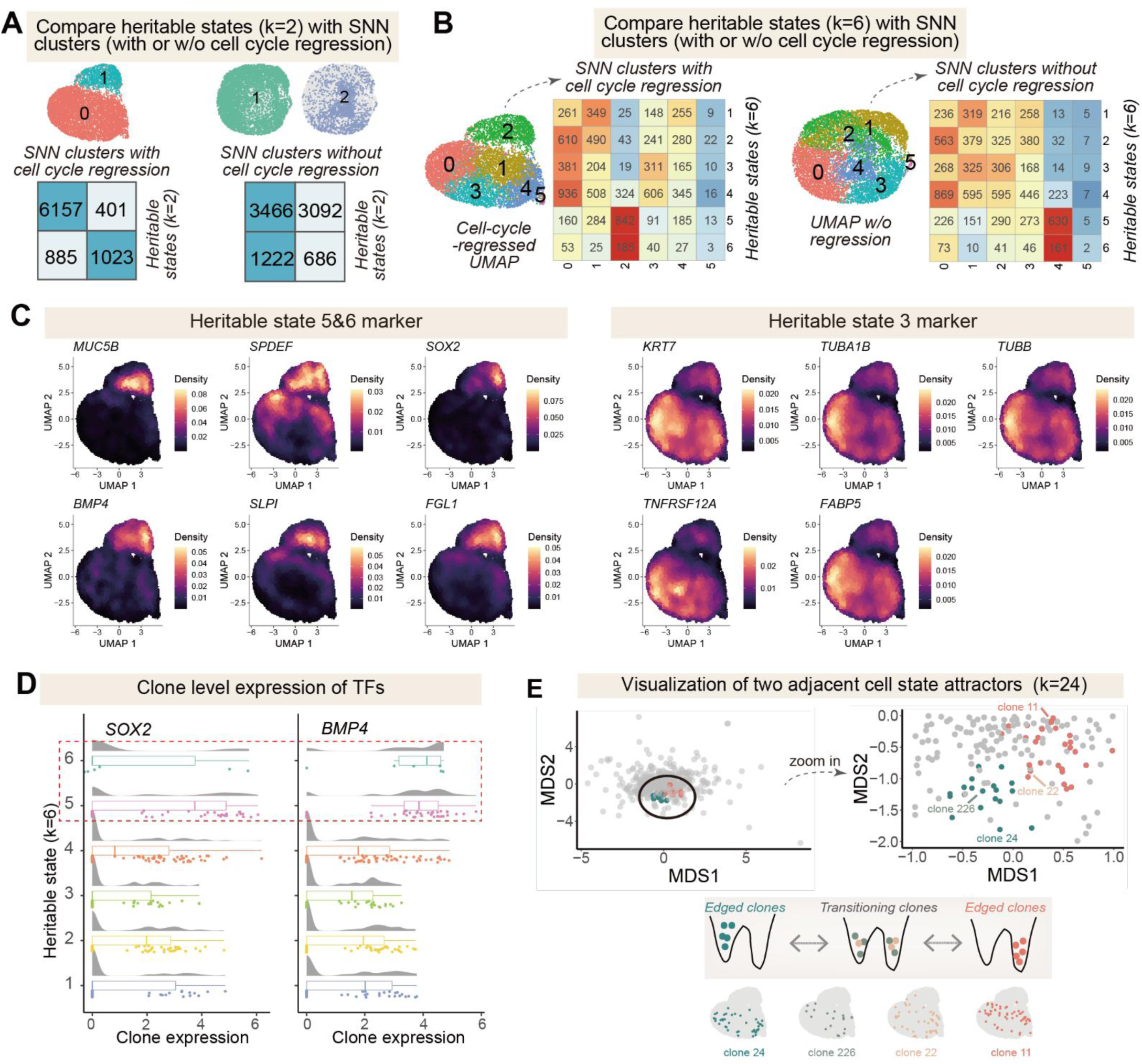
Detailed characterization of heritable states in A549 *in vitro*. **A**, Comparison of heritable states (k=2) with SNN clusters based on single-cell transcriptomes with (left) or without (right) cell cycle regression. Note that the heritable states correspond much better with the SNN clusters based on cell-cycle-regressed single-cell transcriptomes. **B**, Comparison of heritable states (k=6) with SNN clusters based on single-cell transcriptomes with (left) or without (right) cell cycle regression. Confusion matrix showing the correspondence between heritable states and SNN clusters (Seurat, SNN=0.4, k=6). **C**, Density plot of representative heritable state maker genes of A549 cells. These genes may display specific patterns in the UMAP but are not necessarily typical markers of specific transcriptome clusters. **D**, Distribution of *SOX2* and *BMP4* expression at the clone level for each heritable state. Clone-level expression of the largest 200 clones was plotted. Dashed box highlights states of interest. **E**, (top) Visualization of two adjacent heritable states (red: 31 clones, 648 cells; green: 19 clones, 399 cells) in the clonal MDS visualization (k=24). (bottom) Schematics illustrating potential cell state transition between the two states. Some clones are highly localized in one cell state, while others are a mixture of cells from both states. Single-cell UMAP visualizations of clones illustrated were also shown.

**Figure S3:**
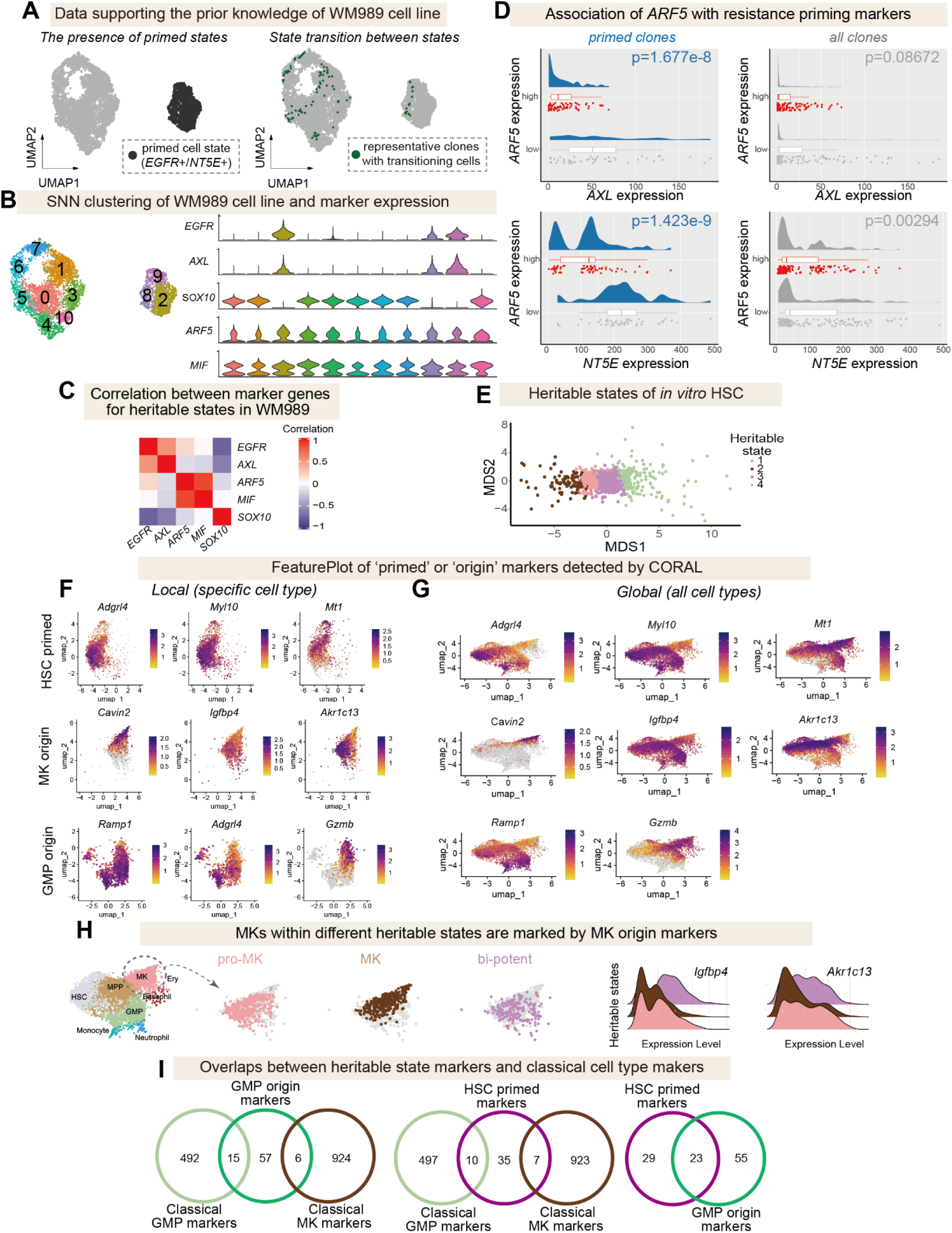
Extended characterization of heritable states in melanoma and HSC cells. **A-D**, Characterization of heritable states in melanoma cells. **A**, UMAP visualization of the *SOX10*+ (sensitive) and *EGFR*+ (primed) states of WM989 cells and clones, showing cells transitioning between states in the UMAP space of a public single-cell lineage tracing dataset^11^. **B**, (left) SNN clustering of the WM989 single-cell RNA-seq dataset as reported in the original publication. (right) Violin plot of key marker genes on the *EGFR* axis and *ARF5* axis along SNN clusters. While the heterogeneity of the *EGFR* axis can be detected by SNN clustering, the heterogeneity of the *ARF5* axis is hidden. **C**, Correlation matrix of marker genes at the clone level in WM989 cells. **D**, Additional characterization for association of *ARF5* level with resistance priming. Within the 122 “primed” clones, *ARF5*-high clones tend to transition to a state with low levels of priming markers (*AXL* or *NT5E*). **E-I**, Characterization of heritable states in mouse HSC cells. **E**, Clonal MDS visualization of HSCs *in vitro* lineage tracing datasets, with clones assigned to specific heritable states labeled by color (pro-MK: 167 clones; MK: 94 clones; bi-potent state: 397 clones, pro-GMP: 183 clones). **F**, Feature plots of primed or origin markers within corresponding cell types (top to bottom: HSC, MK, and GMP), showing heterogeneous expression patterns in single-cell UMAP. These markers cannot be regarded as typical markers of transcriptome clusters representing cell types. **G**, Global feature plots of primed or origin markers in single-cell UMAP, showing distinct patterns along developmental trajectories. **H**, Feature plots and ridgeline plots of representative differentially expressed genes in MK cells from heritable states primed for pro-MK state (state 1), MK state (state 2), and bi-potent state (state 3). **I**, Venn diagram showing the minimal overlap between typical cell type markers and CORAL-detected “primed” or “origin” markers in HSCs.

**Figure S4:**
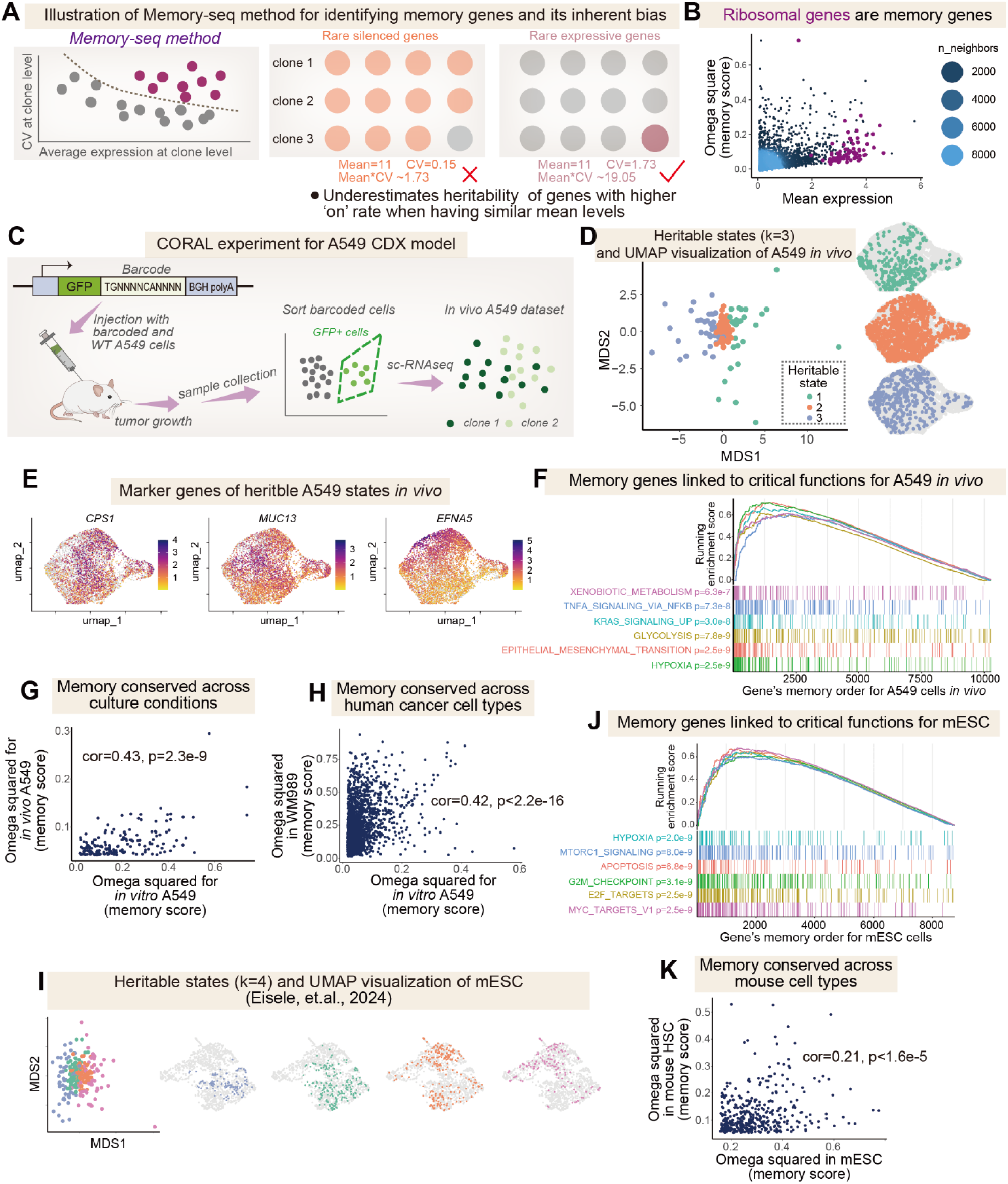
Conservation and functional relevance of transcriptional memory across culture conditions and cell types. **A**, (left) Illustration of Memory-seq method^9^ for memory gene identification. Memory-seq selects genes with relatively high coefficient-of-variation (CV) at the clone level, while our method compares inter- and intra-clone variance. (right) Inherent limitations of the Memory-seq method. Memory-seq method systematically underestimates the heritability of genes with a higher “on” rate, which can lead to biased results in the analysis of gene expression memory Note that yellow and purple genes have the same mean value, but different “on” rates. **B**, Omega squared statistic of all expressed genes in A549 cells, with ribosomal genes (purple) highlighted. **C**, Experimental design for *in vivo* lineage tracing of A549 cells. About 20% of A549 cells transfected with cellular barcodes (green) and 80% wild-type (WT) cells were mixed, and 5 × 10^6^ cells were injected into a mouse to form a CDX (cell-derived xenograft) model. Tumors were allowed to grow for 45 days before isolation, and GFP+ cancer cells were sorted by FACS for single-cell RNA-seq. **D**, Clonal MDS (left) and single-cell UMAP (right) of A549 *in vivo* data. MDS embeddings showing the distribution of 28 clones (green), 58 clones (orange), and 39 clones (blue) assigned to indicated heritable states. UMAP visualization of cells assigned to indicated heritable states: 318 cells (green), 1325 cells (orange), and 415 cells (blue). **E**, Representative marker genes associated with heritable states for A549 *in vivo*. **F**, GSEA analysis using Hallmark gene sets for genes ranked by memory score for A549 *in vivo*. **G**, Correlation between A549 genes’ memory scores (omega squared) *in vitro* and *in vivo*. n=180 shared memory genes. **H**, Correlation between genes’ memory scores in A549 and melanoma cells. n=2593 shared memory genes. **I**, Heritable states (k=4) visualized on clonal MDS (left) and single-cell UMAP (right) for public lineage tracing data of mESCs^51^, with cells within each heritable state relatively localized in UMAP. Green: 71 clones, 197 cells; orange: 83 clones, 299 cells; blue: 44 clones, 115 cells; magenta: 49 clones, 136 cells. **J**, GSEA analysis using Hallmark gene sets for genes ranked by memory score for mESC. **K**, Correlation between genes’ memory scores in mESC and mouse HSC cells. n=403 shared memory genes.

**Figure S5:**
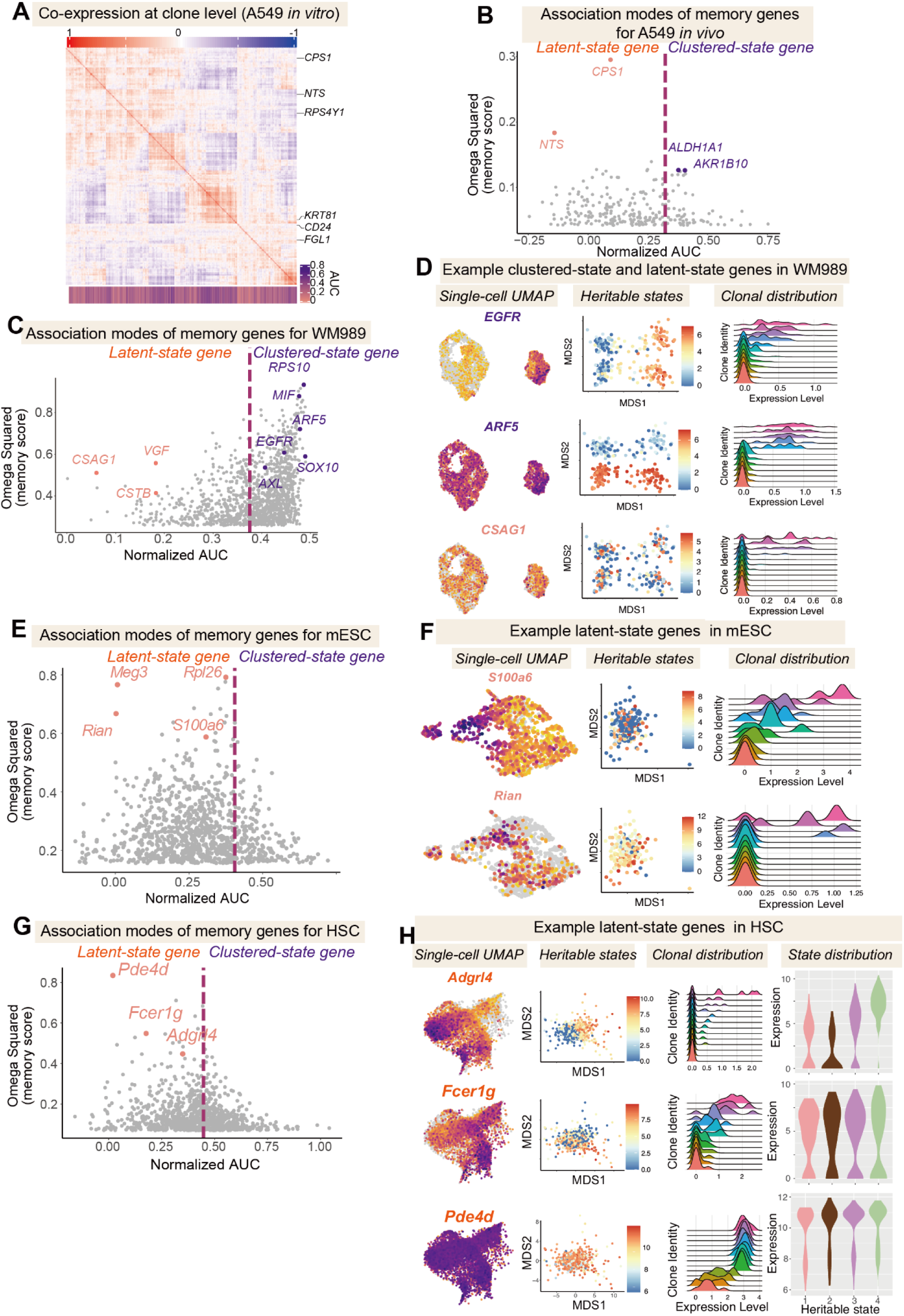
Characterizing association modes of memory genes across cell types. **A**, Co-expression analysis of the top 1000 memory genes in A549 cells *in vitro* at the clone level. Note that clustered-state and latent-state genes are highly correlated. **B**, Quantification of association modes (i.e., AUC) for memory genes (n=252) for A549 cells *in vivo*. Dashed line indicates the global area under the omega squared curve. **C**, Quantification of association modes (i.e., AUC) for top 1000 memory genes in WM989 melanoma cells. Note that there are more clustered-state genes than latent-state genes. **D**, Visualization of representative genes with clustered-state (*EGFR*, *ARF5*) and latent-state (*CSAG1*) genes in melanoma cells, including single-cell UMAP, clonal MDS, and ridgeline plots of clonal gene expression distributions. **E**, Quantification of association modes (i.e., AUC) for top 1000 memory genes in mESC cells. Note that there are more latent-state genes than clustered-state genes. **F**, Visualization of representative memory genes in mESC cells, including single-cell UMAP, clonal MDS, and ridgeline plots of clonal gene expression distributions. **G**, Quantification of association modes (i.e., AUC) for top 1000 memory genes in mouse HSC cells. **H**, Visualization of representative memory genes in HSC cells, including single-cell UMAP, clonal MDS, and ridgeline plots of clonal gene expression distributions.

**Figure S6:**
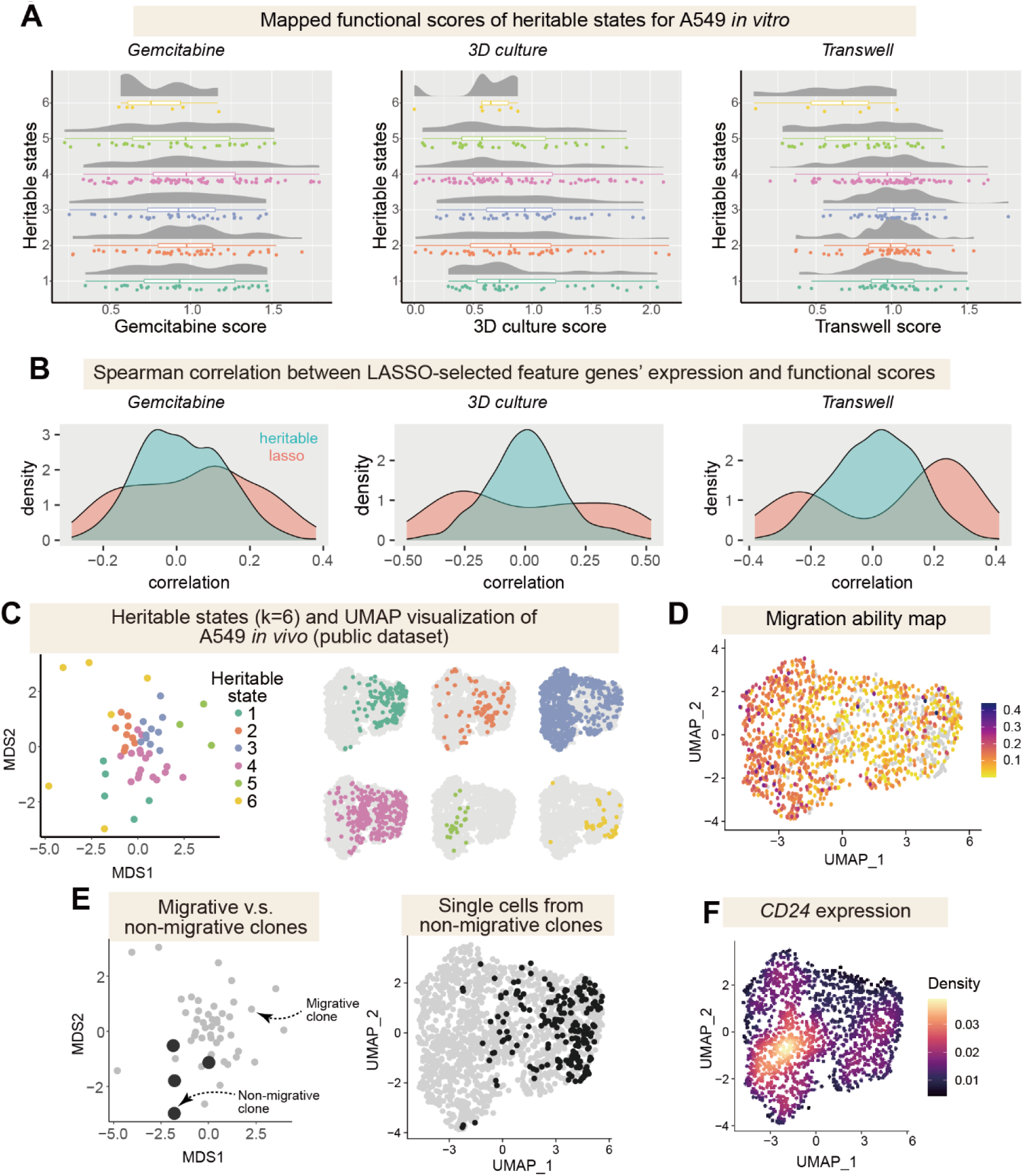
Latent-state memory genes predict cancer-relevant functional phenotypes. **A**, Heritable state-specific distribution of enrichment scores calculated from phenotype mapping experiments for the largest 250 clones, including gemcitabine resistance, 3D culture viability, and Transwell activity. **B**, Distribution of correlation between memory gene expression levels and enrichment scores before and after LASSO regression. The green curves represent the distribution of the top 1000 memory genes, while the red curves represent the distribution of genes selected by LASSO (97 genes associated with gemcitabine resistance, 84 genes associated with 3D culture viability, and 63 genes associated with Transwell activity). **C-F**, Functional characterizations of heritable states using public lineage tracing data of lung cancer metastasis^62^. **C**, Clonal MDS (left) and single-cell UMAP (right) of the largest 50 A549 clones located in the left lung within different heritable states (k=6). **D**, UMAP visualization generated from 1528 cells located in the left lung, labeled with their inferred migration ability. **E**, MDS visualization of non-migrative clones (black, 4 clones) and migrative clones (gray, 46 clones), and location of non-migrative clones in UMAP. Non-migrative clones consist of clonal barcodes not observed in other samples (liver, right lung, etc.). **F**, Density plot of *CD24* expression in single-cell UMAP.

**Figure S7:**
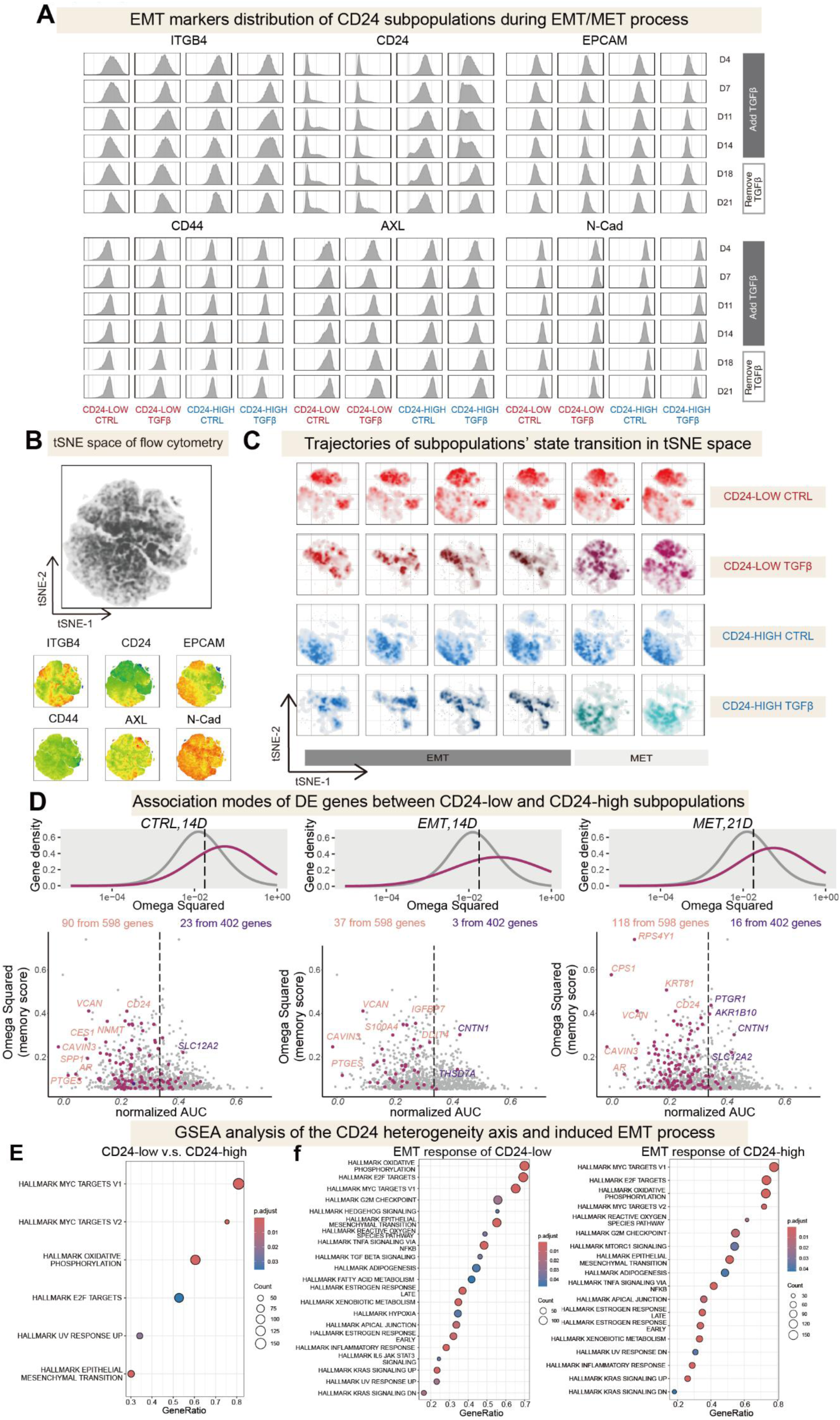
Characterizing the stability of memory genes in A549 cells in response to signaling perturbation. **A**, Flow cytometry measurements of EMT-relevant surface markers in CD24-low and CD24-high A549 subpopulations during the EMT (14 days) and MET (7 days) processes, as well as in the control group. **B**, tSNE embedding of flow cytometry data from all samples, showing the patterns of EMT-relevant surface markers in the tSNE space. **C**, Density plot of CD24-low and CD24-high A549 subpopulations during the EMT and MET processes, measured by flow cytometry. The difference between the CD24-low and CD24-high populations was preserved in the control group but was reduced during the EMT process and recovered during the MET process. **D**, Memory score (i.e., omega squared) and association mode (i.e., AUC) of differentially expressed (DE) genes between CD24-low and CD24-high subpopulations during the EMT/MET process. (top) The gray curve represents the memory score distribution of the top 1000 memory genes, while the purple curve represents the memory score distribution of *CD24*-relevant memory genes. (bottom) In the association mode analysis, the purple dots highlight *CD24*-relevant memory genes within the top 1000 memory genes, most of which are latent-state genes. **E**, GSEA of DE genes between CD24-low and CD24-high subpopulations. Several Hallmark gene sets were enriched, including the epithelial-mesenchymal transition (EMT) gene set. **F**, GSEA of DE genes during the EMT process in CD24-low and CD24-high subpopulations. Several Hallmark gene sets were enriched, including the EMT gene set.

**Figure S8:**
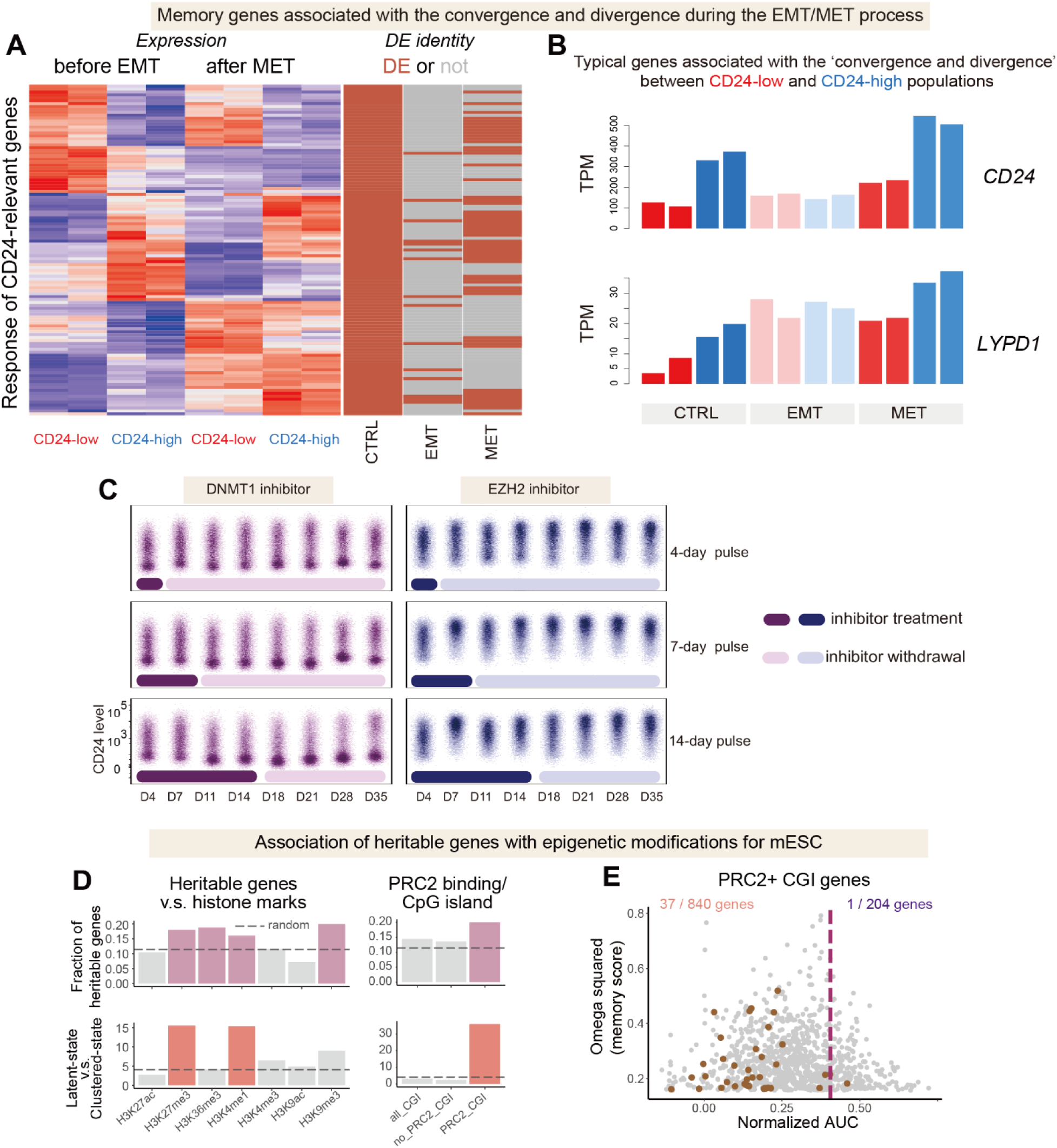
Convergence-and-divergence of memory genes during EMT/MET and their epigenetic maintenance. **A**, Heatmap of DE genes between CD24-low and CD24-high subpopulations before and after the EMT/MET process. The right annotation indicates whether these genes were DE under different conditions (CTRL: 14 days; EMT: 14 days with TGF-β treatment; MET: 21 days with TGF-β withdrawal). Most initial memory DE genes reduce their differences during the EMT process, while some become DE again after the MET process. **B**, Expression levels of representative genes contributing to the “convergence and divergence” between CD24-low (red) and CD24-high (blue) subpopulations during the EMT/MET process. **C**, CD24 expression distribution in A549 cells treated with different pulses of DNMT1 inhibitor (purple) or EZH2 inhibitor (blue), measured by flow cytometry. The up-regulation of CD24 by EZH2 inhibitor and the down-regulation of CD24 by DNMT1 inhibitor persisted even after the withdrawal of the epigenetic inhibitor. **D**, Fraction of memory genes (top) and the ratio between latent-state and clustered-state genes (bottom) in mESC genes with indicated histone modification patterns and PRC2 binding/CGI-enriched patterns. **E**, Association modes of mESC memory genes enriched for PRC2+/CGI (n=38).

## Supplemental Notes

### Supplemental Note 1. Our experimental strategy in relation to existing lineage tracing techniques

Our experimental strategy is similar to previous lineage tracing techniques using static barcodes with high diversity, including LARRY^1^, scMemory-seq^2^ and CellTagging^3^. As the throughput and time resolution of lineage tracing data are limited, each technique has its own trade-off. For example, LARRY, a two-timepoint lineage tracing technique, attempted to maximize the clones detected at both the initial and later timepoints. While such a strategy excels at detecting potential cell transition events, the relatively low cell number per clone limits the quantitative inference of transition kinetics. By contrast, the scMemory-seq technique, captured lineage-encoded transcriptome data at only one timepoint, thus enabling better recovery of each clone’s progeny. However, the initial cell states of clones in the scMemory-seq technique can only be determined by prior FACS enrichment experiment, which can only recover dynamics of several pre-identified genes. The CellTagging technique used a multiple-timepoint barcoding strategy to recover complex lineage relationships in the endpoint snapshot, though the initial state of clones was also not directly observed.

As our work focuses on the lineage-tracing experiment’s potential in detecting hidden “epigenetic properties” to optimize the cell state definition in single-cell experiment, sufficient sampling of clones is crucial (**Figure S1A**). Thus, we chose a relatively simple design in the lineage tracing experiment, similar to scMemory-seq, to maximize the number of cells per clone in one ‘snapshot’ single-cell experiment. This design is easily adaptable to many biological systems, both *in vitro* and *in vivo*. It requires only efficient lentiviral transfection and cell proliferation. We further proposed that such strategy would not limit the power of lineage tracing, even compared with current methods with complex experiment designs^4^. That is, with the CORAL computational workflow designed for snapshot data, single-cell RNA-seq datasets with clone identities can be sufficient for dissecting biological insights into cell heterogeneity and gene expression dynamics.

### Supplemental Note 2. Comparison of CORAL with existing computational methods for lineage analysis

The optimal computational practice for extracting cell state information from lineage tracing data has yet to be established. In most cases, the biological insights were generated from exploratory data analysis from conventional single-cell RNA-seq analysis pipelines, although some computational workflows (e.g., CLiNC^5^, ClonoCluster^6^, LineageOT^7^, Cospar^8^) designed for lineage tracing have been proposed.

A factor limiting the broader application of these pipelines is that few were designed to accommodate diverse experimental designs and biological systems, including even snapshot lineage tracing in cultured cell lines (e.g., scMemory-seq^2^). For example, CLiNC is designed for inference of cell-type hierarchy during development and thus required prior identification of cell types, which can be challenging in highly plastic systems without consensus cell types (e.g., cancer cells with various phenotypes, or the high-resolution analysis of continuous differentiation trajectories). The LineageOT and Cospar algorithms were designed for coarse-graining the time-series lineage tracing data, which generally require at least two timepoints of single-cell RNA-seq data to generate reliable inference. We noted that by estimating the initial states, Cospar may analyze lineage tracing data with only one timepoint. However, it requires a relatively accurate initial “transition map”, which can be inferred from only classical “snapshot” single-cell RNA-seq^9^. Consequently, the cell state identification bias present in the snapshot data may persist even after incorporating lineage information, particularly when the “cell state transition” landscape within the snapshot is unclear.

In other words, most methods were still highly reliant on the cell state information from prior biological knowledge or from conventional analyses of single-cell RNA-seq data. This contradicts a key motivation of lineage tracing: to accurately infer the epigenetic landscape (i.e., cell state divisions and transition kinetics), as relying on single-cell RNA-seq data alone can introduce distortions. To overcome the challenges, the existence of hidden variables needs to be modeled explicitly in the beginning of the workflow, with the help of clone identities. That is, most hidden variables (also known as epigenetic states compared with the transcriptome states) were shared within a clone. In fact, the ClonoCluster method is an inspiring attempt to introduce clone information in the pre-processing process, including UMAP visualization and clustering. However, the “weight” of the additional clone information in ClonoCluster was only determined empirically, making it more of an exploratory tool than a method for quantitative inference.

By contrast, the CORAL computational workflow systematically addresses these limitations. Firstly, CORAL was designed for the simplest case of lineage tracing (i.e., the single-cell RNA-seq data with initial clone identities), which can be easily collected with acceptable cost. Existing lineage tracing datasets with complex designs can be divided into separate “snapshot” datasets for analysis with CORAL. Yet, interpreting multiple CORAL results requires further analysis. Secondly, CORAL demonstrated good performance in both steady-state cultured cell lines and non-steady-state developmental processes, especially for dissecting cell states not visible in conventional analysis. Thirdly, CORAL’s workflow is orthogonal to current single-cell RNA-seq pre-processing workflows, thus preventing bias introduced by feature selection, dimensionality reduction, and clustering.

CORAL demonstrates clear theoretical motivations and guidance for integrating the clonal and transcriptional information. Our premise is that the significant transcriptional heterogeneity between clones, rather than within them, accurately reflects the energetic “barriers” between cell states and should thus be the basis for defining heritable cell states, or micro-attractors. In other words, clusters of clones with similar transcriptome distributions, rather than just clusters of cells, indicate metastable micro-attractors, or valleys with specific transcriptional (explicit) and epigenetic (hidden) features. Such motivation is, in essence, another way of describing the assumption of CLiNC^5^, which suggests that the cell types with shared clonal barcodes detected should be closer in the lineage hierarchy tree. However, we expand the idea to the initial definition of “cell states”, making full use of the transcriptome measurements, instead of just using prior biological knowledge.

Expanding the CORAL modeling framework to infer the cell state transition map (like in Cospar and lineageOT) would be desirable. However, modeling both transcriptional and epigenetic cell state transition can be challenging, especially for the highly plastic biological systems (e.g., cancer cells or HSCs from mouse bone marrow). In such situations, most previous cell state transition inference methods have performed poorly, often failing to identify reliable transcriptional and epigenetic cell states. As for the CORAL framework, though it can outperform other methods in the reconstruction of cell state structures in those systems, several key issues still need to be solved: 1) Estimating accurate initial cell states when transcriptome-based inference (e.g., WaddingtonOT^9^) is unreliable, 2) accurately defining the state of each individual cell. We expect that with the increasing number of well-sampled “ground truth” datasets with multiple timepoints and multi-omics measurements, CORAL can be expanded to predict more cell state transition kinetics than its current version.

### Supplemental Note 3. The theoretical basis of CORAL

From the perspective of dynamical systems, the quantitative modeling of single-cell omics data typically includes the following steps: 1) the definition of dominant attractors in the landscape (i.e., cell types/cell states), 2) reconstruction of the vector field (i.e., the developmental/noisy trajectories of cells), and 3) the inference of the parameters of dynamical system (e.g., the regulatory relationships)^10^.

Current approaches to analyzing single-cell data often use the distribution of cells in high-dimensional gene expression space to infer properties of the underlying dynamical system. These approaches are based on the decomposition of the Fokker-Planck equation according to potential landscape theory^11^ and quasi-potential theory^12,13^. To clarify the theoretical foundation of CORAL, we adapt the following equations from established literatures^12,13^.

The Fokker-Planck equation, which describes the time evolution of the probability distribution *P*(*x*, *tt*) of cells in state space x, is given by:

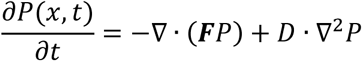

where ***F*** represents the deterministic force field and *D* is the diffusion matrix representing noise.

Under potential landscape theory, the dynamics can be expressed as the gradient of potential function

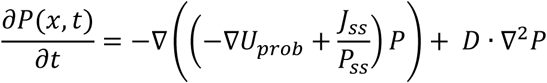

where ∇*U*_*p*_*_rob_* = −∇(− ln(*P*_*ss*_)) · *D*, and *P*_*ss*_ represents the steady-state distribution.

Quasi-potential theory introduces the concept of a global quasi-potential, *U_global-quasi_*, related to the Freidlin-Wentzell action, which characterizes transition rates between stable states along the least action path (LAP). The LAP is the most probable trajectory for a system to transition between two stable states in the presence of noise. The dynamics under this theory can be written as:

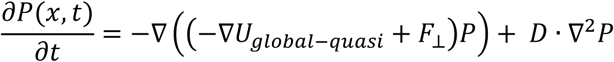

where (∇*U_global_*_−*quasi*_, *F*_⊥_) = 0, indicating that *F*_⊥_ is orthogonal to the gradient of the global quasi-potential.

Notably, *U_prob_* equals *U_global-quasi_* when (−∇*U_prob_*, 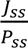 = 0. In other words, it is *U_global_*_−quasi_, not *U_prob_*, that is mathematically linked to the Freidlin-Wentzell potential (and thus to the local quasi-potential, *U_local_*_−*quasi*_), where the spontaneous transition rate is proportional to *e*^−*U_local−quasi_*^. Intuitively, the global and local quasi-potentials can be related through a “pruning and sticking” procedure^13^.

Those properties of dynamical systems present challenges for Waddington landscape reconstruction from biological snapshot data. Firstly, in a highly plastic landscape, there will be a divergence item in 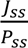 (e.g., cell cycle). Then, the potential landscape inferred from snapshot measurement does not necessarily equal the global quasi-potential and cannot be directly related to the cell state transition rate. Secondly, estimating the cell state transition rate (i.e., calculating local quasi-potential from global quasi-potential) requires knowledge of the saddle point properties^14^. However, for some rare cell state transition events, the transitioning cells are hard to sample^2^.

Thus, a key motivation of CORAL is to leverage lineage tracing data to explore the Waddington landscape using an alternative strategy: searching from the micro-attractor (denoting sub-state of the cell) to obtain information of local quasi-potential, and then “merge” local quasi-potentials to generate a global understanding.

Traditional methods for approximating the global quasi-potential rely on steady-state measurement under conditions of fixed, low noise and sufficient waiting time^14^, as described by:

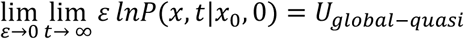

to approximate the global quasi-potential. In contrast, CORAL uses the distribution of a clone originating from a specific cell with state *x*_0_, observed after a fixed time under conditions of assumed low noise^14^, to approximate the local quasi-potential:

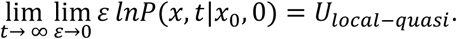

This approximation rests on the concept that progeny cells, confined to a micro-attractor’s basin, explore this local region and exhibit a distribution shaped by the “local” quasi-potential landscape at intermediate timescales. As time progresses, this distribution could approach the global quasi-potential landscape.

This same rationale can explain why lineage tracing data, even from a single timepoint, provides more insight into cell state transition kinetics than typical single-cell RNA-seq data^1,8^, as it is not influenced by non-equilibrium steady states or the lack of transitioning cells^2^. However, the lineage tracing data is always sparser and noisier^8^ due to the temporal nature of the data so the pruning and sticking procedure is not trivial. Traditional clustering or coarse-graining methods, usually still using the high-density clusters in snapshot data, might not be accurate due to hidden states or fluctuations at various timescales.

In the following discussion, we will further propose how the CORAL workflow, which better makes use of the clone identities in the beginning of cell state definition or coarse-graining, further improves our understanding of heritable cell states.

In addition to typical assumptions in Waddington landscape modeling (e.g., epigenetic/transcriptomic cell states are heritable), CORAL assumes that clones with similar distributions in high-dimensional space originate from “initial cells” residing within the same micro-attractor. This allows us to cluster these clones as representatives of specific cell state micro-attractors.

To better approximate this assumption, we used energy distance^15^ to measure the similarity between clones. As the energy distance can be written as:

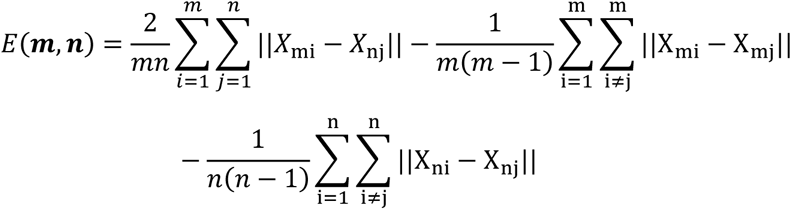

where *m*, *n* denote clone identities and *ii*, *j* are cell identities within corresponding clones.

We noticed that such distance measures smooth the “fast and noisy” axis to maximize the effect of variables that really matter in the local quasi-potential. That is, because noisy genes exhibit heterogeneity within clones, the magnitude of inter-clone distances diminishes the effect of intra-clone distances. Therefore, if two initial cells originate from two unconnected micro-attractors, the energy distance between them will be relatively large, even if the initial Euclidean distance between the micro-attractors in the original transcriptome space is small or even undetectable (i.e., in the case of a hidden epigenetic state). In other words, the energy distance calculation between clones inherently performs feature selection, prioritizing highly heritable features for downstream analysis.

Furthermore, the agglomerative hierarchical clustering using Ward’s method^16^ in CORAL is based on the principle that connected micro-attractors will also be connected in the space defined by the energy distance. For example, if one clone starts from micro-attractor A and has 30% of cells at micro-attractor B due to the A→B transition activity, this clone would be located just between the “all-A” clones and “all-B” clones in the MDS visualization^17^ of energy distance. Because the density of these “transitioning” clones depends on the plasticity between micro-attractors, micro-attractors separated by lower energy barriers are more likely to merge first.

## Supplemental Tables

**Table S1.**
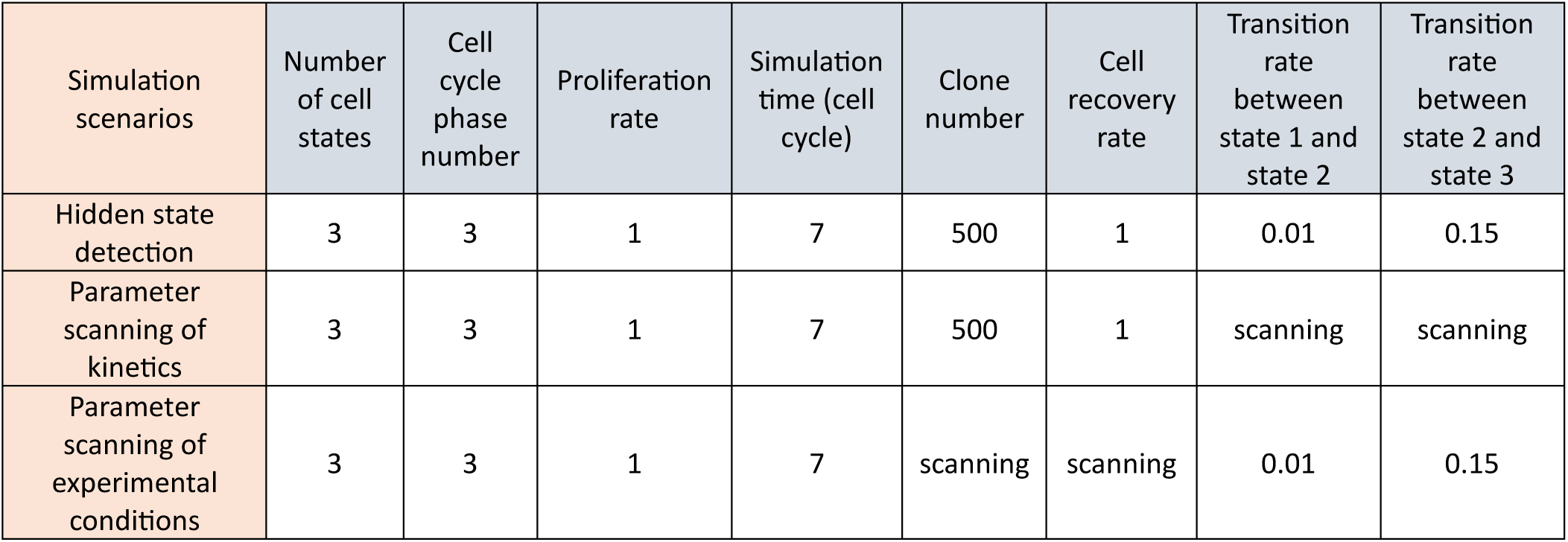
Parameter settings for CORAL simulations.

**Table S2. Heritable cell state (k=6) markers for A549 *in vitro* dataset (see the excel file).**

**Table S3. Marker genes for HSC dataset (see the excel file).**

**Table S4. Memory gene list for A549 *in vitro* dataset (see the excel file).**

**Table S5. Memory gene list for A549 *in vivo* dataset (see the excel file).**

**Table S6. Memory gene list for WM989 melanoma dataset (see the excel file).**

**Table S7. Memory gene list for HSC dataset (see the excel file).**

**Table S8. Memory gene list for mESC dataset (see the excel file).**

**Table S9. Conservation of memory genes across datasets (see the excel file).**

**Table S10. Differentially expressed genes related to Figure 6 (see the excel file).**

**Table S11.**
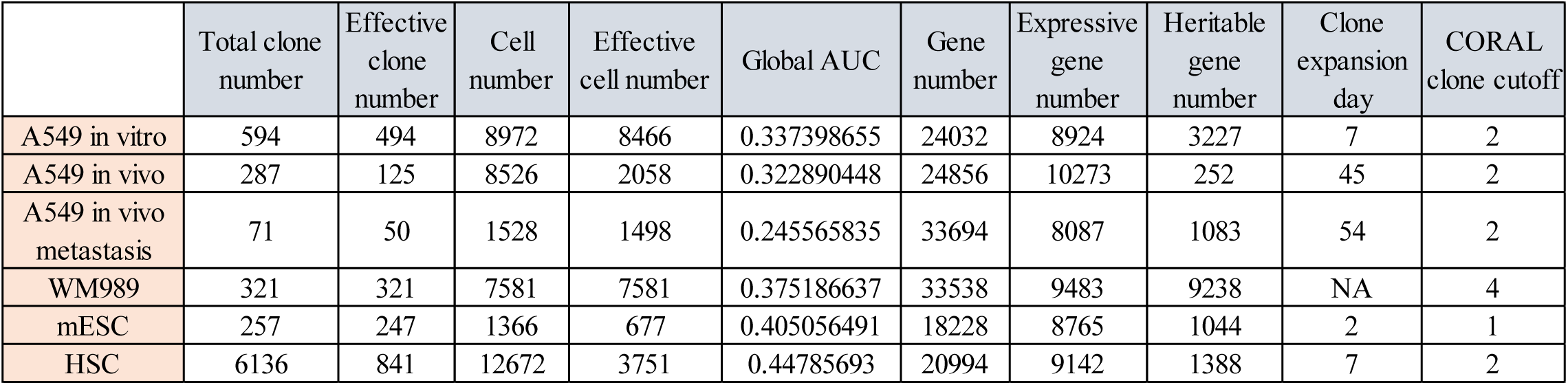
Statistics of all datasets used for CORAL analysis.

## Notes

https://drive.google.com/drive/folders/1-cNiSKZFyVSs9Mndq87AcRXfaGweLesj?usp=sharing.

### Competing Interest Statement

The authors have declared no competing interest.

