## Supplemental Notes for "Widespread transcriptional memory shapes heritable states and functional heterogeneity in cancer and stem cells"

The Fokker-Planck equation, which describes the time evolution of the probability distribution $P(x,t)$ of cells in state space x, is given by:

$$\frac{\partial P(x,t)}{\partial t} = -\nabla \cdot (\boldsymbol{F}P) + D \cdot \nabla^2 P$$

where $\boldsymbol{F}$ represents the deterministic force field and $D$ is the diffusion matrix representing noise.

Under potential landscape theory, the dynamics can be expressed as the gradient of a potential function $U_{prob}$:

$$\frac{\partial P(x,t)}{\partial t} = -\nabla\left(\left(-\nabla U_{prob} + \frac{J_{ss}}{P_{ss}}\right)P\right) + D \cdot \nabla^2 P$$

where $\nabla U_{prob} = -\nabla(-\ln(P_{ss})) \cdot D$, and $P_{ss}$ represents the steady-state distribution.

Quasi-potential theory introduces the concept of a global quasi-potential, $U_{global-quasi}$, related to the Freidlin-Wentzell action, which characterizes transition rates between stable states along the least action path (LAP). The LAP is the most probable trajectory for a system to transition between two stable states in the presence of noise. The dynamics under this theory can be written as:

$$\frac{\partial P(x,t)}{\partial t} = -\nabla\left(\left(-\nabla U_{global-quasi} + F_{\perp}\right)P\right) + D \cdot \nabla^2 P$$

where $\left(\nabla U_{global-quasi}, F_{\perp}\right) = 0$, indicating that $F_{\perp}$ is orthogonal to the gradient of the global quasi-potential.

Notably, $U_{prob}$ equals $U_{global-quasi}$ when $\left(-\nabla U_{prob}, \frac{J_{ss}}{P_{ss}}\right) = 0$ . In other words, it is $U_{global-quasi}$, not $U_{prob}$, that is mathematically linked to the Freidlin-Wentzell potential (and thus to the local quasi-potential, $U_{local-quasi}$), where the spontaneous transition rate is proportional to $e^{-U_{local-quasi}}$. Intuitively, the global and local quasi-potentials can be related through a "pruning and sticking" procedure[13].

Those properties of dynamical systems present challenges for Waddington landscape reconstruction from biological snapshot data. Firstly, in a highly plastic landscape, there will be a divergence item in $\frac{J_{ss}}{P_{ss}}$ (e.g., cell cycle). Then, the potential landscape inferred from snapshot measurement does not necessarily equal the global quasi-potential and cannot be directly related to the cell state transition rate. Secondly, estimating the cell state transition rate (i.e., calculating local quasi-potential from global quasi-potential) requires knowledge of the saddle point properties[14]. However, for some rare cell state transition events, the transitioning cells are hard to sample[2].

Thus, a key motivation of CORAL is to leverage lineage tracing data to explore the Waddington landscape using an alternative strategy: searching from the micro-attractor (denoting sub-state of the cell) to obtain information of local quasi-potential, and then "merge" local quasi-potentials to generate a global understanding.

Traditional methods for approximating the global quasi-potential rely on steady-state measurement under conditions of fixed, low noise and sufficient waiting time[14], as described by:

$$\lim_{\varepsilon \to 0} \lim_{t \to \infty} \varepsilon \, lnP(x, t | x_0, 0) = U_{global-quasi}$$

to approximate the global quasi-potential. In contrast, CORAL uses the distribution of a clone originating from a specific cell with state $x_0$, observed after a fixed time under conditions of assumed low noise[14], to approximate the local quasi-potential:

$$\lim_{t \to \infty} \lim_{\varepsilon \to 0} \varepsilon \, lnP(x, t | x_0, 0) = U_{local-quasi}.$$

This approximation rests on the concept that progeny cells, confined to a micro-attractor's basin, explore this local region and exhibit a distribution shaped by the "local" quasi-potential landscape at intermediate timescales. As time progresses, this distribution could approach the global quasi-potential landscape.

This same rationale can explain why lineage tracing data, even from a single timepoint, provides more insight into cell state transition kinetics than typical single-cell RNA-seq data[1,8], as it is not influenced by non-equilibrium steady states or the lack of transitioning cells[2]. However, the lineage tracing data is always sparser and noisier[8] due to the temporal nature of the data so the pruning and sticking procedure is not trivial. Traditional clustering or coarse-graining methods, usually still using the high-density clusters in snapshot data, might not be accurate due to hidden states or fluctuations at various timescales.

In the following discussion, we will further propose how the CORAL workflow, which better makes use of the clone identities in the beginning of cell state definition or coarse-graining, further improves our understanding of heritable cell states.

In addition to typical assumptions in Waddington landscape modeling (e.g., epigenetic/transcriptomic cell states are heritable), CORAL assumes that clones with similar distributions in high-dimensional space originate from "initial cells" residing within the same micro-attractor. This allows us to cluster these clones as representatives of specific cell state micro-attractors.

To better approximate this assumption, we used energy distance[15] to measure the similarity between clones. As the energy distance can be written as:

$$E(\boldsymbol{m},\boldsymbol{n}) = \frac{2}{mn}\sum_{i=1}^{m}\sum_{j=1}^{n}||X_{\mathrm{mi}} - X_{\mathrm{nj}}|| - \frac{1}{m(m-1)}\sum_{\mathrm{i=1}}^{\mathrm{m}}\sum_{\mathrm{i\neq j}}^{\mathrm{m}}||\mathrm{X_{mi}} - \mathrm{X_{mj}}|| - \frac{1}{n(n-1)}\sum_{\mathrm{i=1}}^{\mathrm{n}}\sum_{\mathrm{i\neq j}}^{\mathrm{n}}||\mathrm{X_{ni}} - \mathrm{X_{nj}}||$$

where $m, n$ denote clone identities and $i, j$ are cell identities within corresponding clones.

We noticed that such distance measures smooth the "fast and noisy" axis to maximize the effect of variables that really matter in the local quasi-potential. That is, because noisy genes exhibit heterogeneity within clones, the magnitude of inter-clone distances diminishes the effect of intra-clone distances. Therefore, if two initial cells originate from two unconnected micro-attractors, the energy distance between them will be relatively large, even if the initial Euclidean distance between the micro-attractors in the original transcriptome space is small or even undetectable (i.e., in the case of a hidden epigenetic state). In other words, the energy distance calculation between clones inherently performs feature selection, prioritizing highly heritable features for downstream analysis.

Furthermore, the agglomerative hierarchical clustering using Ward's method[16] in CORAL is based on the principle that connected micro-attractors will also be connected in the space defined by the energy distance. For example, if one clone starts from micro-attractor A and has 30% of cells at micro-attractor B due to the A→B transition activity, this clone would be located just between the “all-A” clones and “all-B” clones in the MDS visualization[17] of energy distance. Because the density of these “transitioning” clones depends on the plasticity between micro-attractors, micro-attractors separated by lower energy barriers are more likely to merge first.

### Supplemental Tables

**Table S1. Parameter settings for CORAL simulations.**

| Simulation scenarios | Number of cell states | Cell cycle phase number | Proliferation rate | Simulation time (cell cycle) | Clone number | Cell recovery rate | Transition rate between state 1 and state 2 | Transition rate between state 2 and state 3 |
|---|---|---|---|---|---|---|---|---|
| Hidden state detection | 3 | 3 | 1 | 7 | 500 | 1 | 0.01 | 0.15 |
| Parameter scanning of kinetics | 3 | 3 | 1 | 7 | 500 | 1 | scanning | scanning |
| Parameter scanning of experimental conditions | 3 | 3 | 1 | 7 | scanning | scanning | 0.01 | 0.15 |

**Table S2. Heritable cell state (k=6) markers for A549 *in vitro* dataset (see the excel file).**

**Table S3. Marker genes for HSC dataset (see the excel file).**

**Table S4. Memory gene list for A549 *in vitro* dataset (see the excel file).**

**Table S5. Memory gene list for A549 *in vivo* dataset (see the excel file).**

**Table S6. Memory gene list for WM989 melanoma dataset (see the excel file).**

**Table S7. Memory gene list for HSC dataset (see the excel file).**

**Table S8. Memory gene list for mESC dataset (see the excel file).**

**Table S9. Conservation of memory genes across datasets (see the excel file).**

**Table S10. Differentially expressed genes related to Figure 6 (see the excel file).**

**Table S11. Statistics of all datasets used for CORAL analysis.**

| | Total clone number | Effective clone number | Cell number | Effective cell number | Global AUC | Gene number | Expressive gene number | Heritable gene number | Clone expansion day | CORAL clone cutoff |
|---|---|---|---|---|---|---|---|---|---|---|
| A549 in vitro | 594 | 494 | 8972 | 8466 | 0.337398655 | 24032 | 8924 | 3227 | 7 | 2 |
| A549 in vivo | 287 | 125 | 8526 | 2058 | 0.322890448 | 24856 | 10273 | 252 | 45 | 2 |
| A549 in vivo metastasis | 71 | 50 | 1528 | 1498 | 0.245565835 | 33694 | 8087 | 1083 | 54 | 2 |
| WM989 | 321 | 321 | 7581 | 7581 | 0.375186637 | 33538 | 9483 | 9238 | NA | 4 |
| mESC | 257 | 247 | 1366 | 677 | 0.405056491 | 18228 | 8765 | 1044 | 2 | 1 |
| HSC | 6136 | 841 | 12672 | 3751 | 0.44785693 | 20994 | 9142 | 1388 | 7 | 2 |
